# Drosophila RASopathy Models Identify Disease Subtype Differences and Biomarkers of Drug Efficacy

**DOI:** 10.1101/2020.10.30.362350

**Authors:** Tirtha K. Das, Jared Gatto, Rupa Mirmira, Ethan Hourizadeh, Dalia Kaufman, Bruce D. Gelb, Ross Cagan

**Affiliations:** Dept. of Cell, Developmental, and Regenerative Biology, Icahn School of Medicine at Mount Sinai, New York NY USA; The Mindich Child Health and Development Institute, Dept. of Pediatrics, Department of Genetics and Genomic Science, Icahn School of Medicine at Mount Sinai, New York NY USA; Institute of Cancer Sciences at University of Glasgow, Wolfson Wohl Cancer Research Centre, Glasgow Scotland UK

## Abstract

RASopathies represent a family of mostly autosomal dominant diseases that are caused by missense variants in the RAS/MAPK pathway. In aggregate, they are among the more common Mendelian disorders. They share overlapping pathologies that include structural birth and developmental defects that affect the heart, craniofacial and skeletal, lymphatic, and nervous systems. Variants in different genes—including those encoding KRAS, NRAS, BRAF, RAF1, and SHP2—are associated with overlapping but distinct phenotypes. Here, we report an analysis of 13 Drosophila transgenic lines, each expressing a different human disease isoform associated with a form of RASopathy. Similar to their human counterparts, each Drosophila line has common aspects but also important phenotypic distinctions including signaling pathways as well as response to therapeutics. For some lines, these differences represent activation of pathways outside the core RAS signaling pathway including the Hippo and SAPK/JNK signaling networks. We identified two classes of clinically relevant drugs, statins and histone deacetylase inhibitors, that improved viability across most RASopathy lines; in contrast, several canonical RAS pathway inhibitors proved poorly effective against, *e.g.*, SHP2-expressing lines encoded by *PTPN11*. Our study provides a whole animal platform for comparison of a large number of RASopathy-associated variants. Among these variants we have identified differences in tissue phenotypes, in activation signaling pathways in biomarkers of disease progression and drug efficacy, and suggest drug classes that can be tolerated over long treatment periods for consideration in broad RASopathy trials.

## Introduction

RASopathies are a family of Mendelian diseases defined by germline variants that generally elevate activity of the RAS signaling pathway (Jindal et al., 2017, 2015; Rauen, 2013). RASopathy phenotypes are pleiomorphic with structural birth defects altering the cardiovascular, craniofacial, skeletal, lymphatic, and central nervous systems in addition to postnatal short stature and developmental delays. Found in perhaps 1:1000 new births, RASopathies are associated with variants in genes encoding multiple RAS pathway components including KRAS, NRAS, BRAF, RAF1, and SHP2. SHP2, encoded by the gene *PTPN11*, is a cytosolic protein that belongs to the protein tyrosine phosphatase superfamily, Following growth factor stimulation, SHP2 is recruited to the intracellular domain of receptor tyrosine kinases (RTKs) to activate the RAS/RAF/MAPK signal transduction pathway (Liu and Qu, 2011). Regarding RAS pathway inhibitors as therapies, a recent publication reported some success with trametinib for *RIT1*-associated Noonan syndrome infants with severe hypertrophic cardiomyopathy (Andelfinger et al., 2019). Overall, however, most patients with RASopathy have limited therapeutic options.

One of the challenges in understanding and developing therapeutics for RASopathy patients is the heterogeneity associated with different mutational variants (Rauen, 2013). Noonan syndrome (NS), NS with multiple lentigines (NSML), and NS-like syndromes are associated with variants in multiple genes including *PTPN11*, *KRAS*, *NRAS*, and *RAF1*. Costello syndrome is associated with variants in *HRAS*, while cardiofaciocutaneous (CFC) syndrome is associated with variants in *BRAF*, *MEK1*, *MEK2*, and *KRAS*. These differences are associated with important differences in disease presentation (Rauen, 2013). For example, regarding heart defects, approximately 20% of patients with NS report hypertrophic cardiomyopathy (HCM), a number that increases to 80-90% in NSML patients; young infants with NS-associated HCM have a one-year survival rate of only 34%(Wilkinson et al., 2012). Genotype-phenotype correlations also exist at the gene level: patients with NS due to *RAF1* variants show high prevalence of HCM while *PTPN11* variants are negatively associated with HCM (Pandit et al., 2007; Tartaglia et al., 2002).

We currently have a poor understanding of the mechanisms by which different RASopathy-associated variants—each directing elevated RAS pathway activity—can lead to different patient outcomes. A systematic animal model comparison across a large cross-section of the different RASopathy variants has not been performed: rare Mendelian diseases can prove challenging for targeting specific disease isoforms in clinical trials, and identifying common therapeutic strategies across variants can help address this issue. Further, treatments will need to be well tolerated for extended periods of time, emphasizing a key advantage of whole animal models.

Here, we develop and characterize 13 Drosophila RASopathy models, developing a platform that takes a broad, whole animal approach to exploring the signaling differences and therapeutic responses between RASopathy isoforms. Expressing human transgenes in Drosophila epithelia, we demonstrate significant whole animal signaling differences between RASopathy-associated genes and also between models containing variants within the same gene. These differences include tissue-specific phenotypes as well as activation of different signaling networks within the same tissue. Finally, we observe significant differences in response to clinical drugs that inhibit diverse cellular targets as well as tool compounds that inhibit components of the mitogen activated protein kinase (MAPK) pathway. Overall, these results emphasize differences between disease isoforms that may impact disease progression as well as therapeutics. Nevertheless, we demonstrate at least two classes of drugs that provide broad therapeutic benefits across different RASopathy models, suggesting they have the potential to benefit a large cross-section of patient classes in the clinic.

## Results

Previous work reported Drosophila RASopathy models in which fly genes—altered to model human RASopathy disease isoforms—were expressed in multiple tissue types (Cordeddu et al., 2015; Oishi et al., 2009, 2006). The resulting phenotypes generally emulated RAS gain-of-function phenotypes. In this study, we have generated 13 new Drosophila RASopathy models. Each line expresses a human transgene containing a commonly observed RASopathy variant; each transgene is under the control of a GAL4-inducible UAS promoter (Brand and Perrimon, 1993). We then used this set of 13 lines to explore how the different human transgenes directed similar or different fly phenotypes, altered cellular signaling networks, and responded to drugs. Signaling paradigms uncovered in select RASopathy models were further evaluated using genetic modifier tests. Figure 1A summarizes our approach.

**Figure 1.**
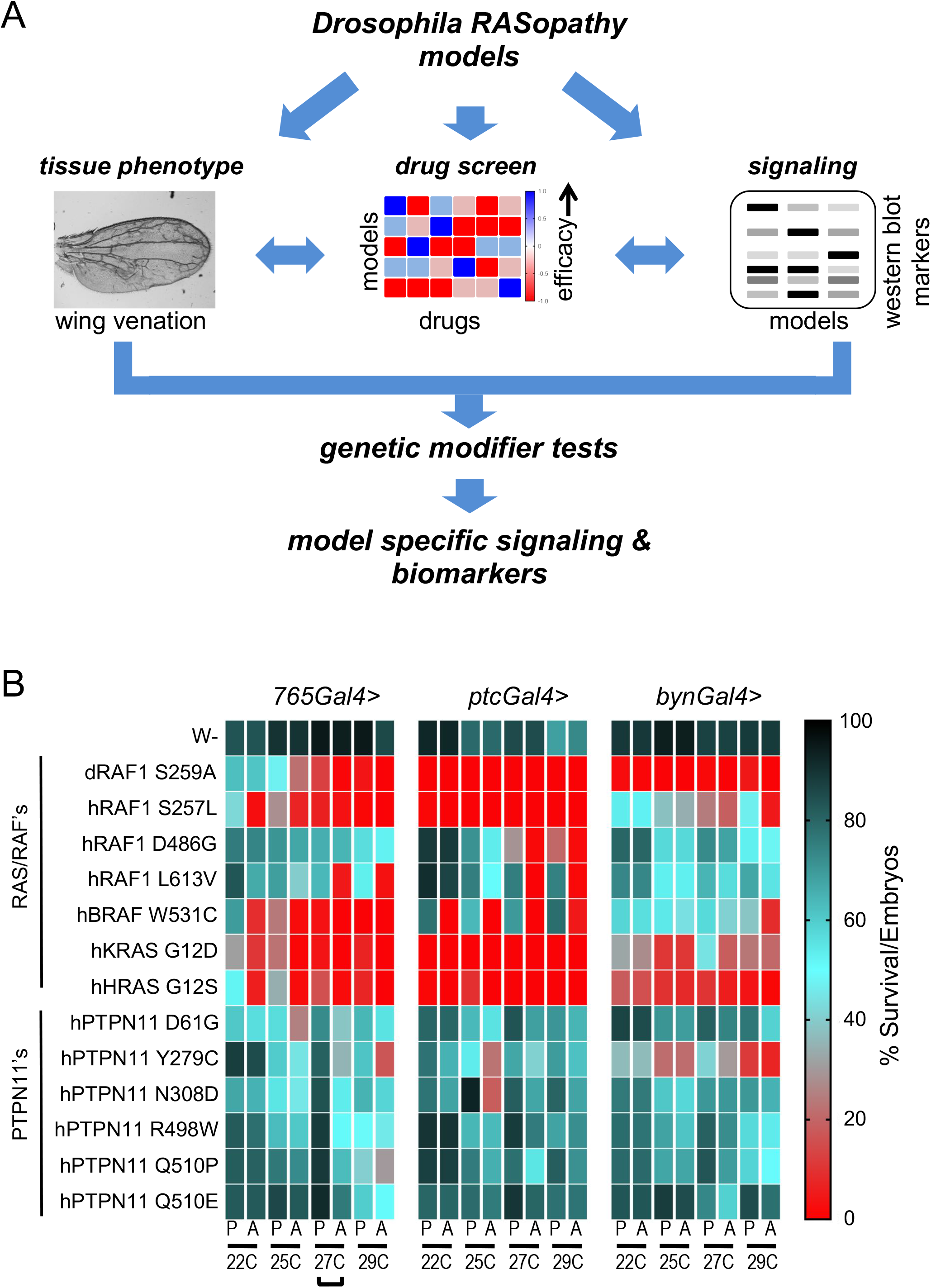
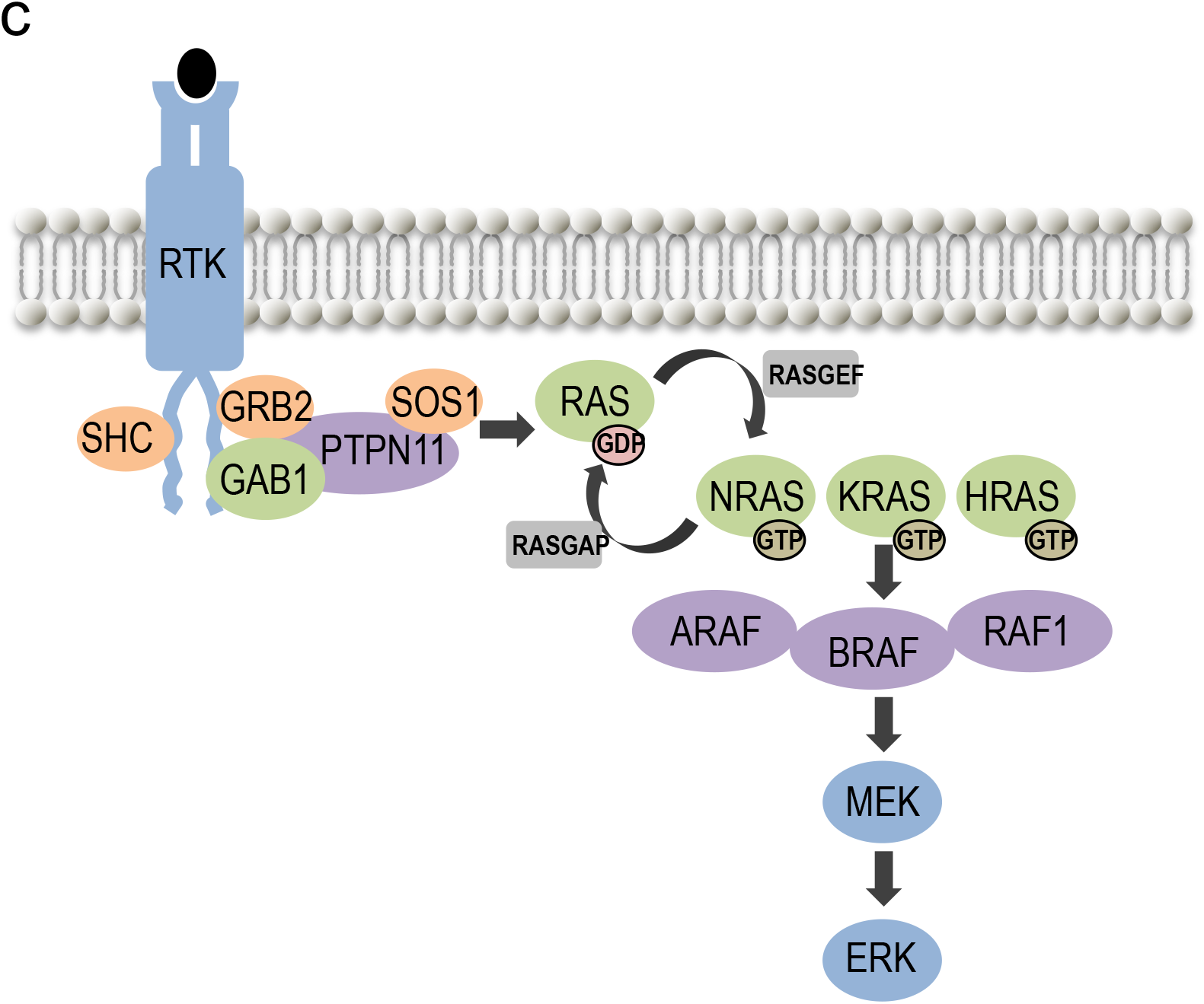
(A) A schematic of the approach, using Drosophila models that expressed different human RASopathy isoforms. Each isoform was induced in flies using different tissue-specific promoters and its effect on wing phenotype assessed (tissue phenotype). Flies were subjected to whole animal screening to identify optimal therapeutics for each model (drug screen). Differences in signaling among isoforms were assessed using western blot analysis in the presence or absence of identified drug hits (signaling). For select models, pathways identified using this approach were functionally validated through genetic knockdown experiments. This integrated approach provided a broad overview of differences in signaling among isoforms and potential biomarkers of therapeutic efficacy. (B) Quantitative viability assay. GAL4 levels progressively increase with increasing temperature, which results in increased transgene expression. The result was increasing lethality, allowing identification of optimal lethality conditions. Percent viability represents the number of pupae (P) or eclosed adults (A) after 12–14 days divided by the total number of embryos originally present in each experiment and is depicted as a heat map. Percent viability color code is shown next to the heat map. The different GAL4 drivers tested in this assay and their primary domain of expression are *765-GAL4* (entire wing), *ptc-GAL4* (several tissues including the central portion of wing), and *byn-GAL4* (hindgut). Bracket highlights the optimal screening condition (*765-GAL4* at 27 °C) that was used for drug screening (Figure 3). This heat map represents approximately 32,000 screened embryos. (C) RASopathy-relevant genes define important aspect of the RAS pathway. *Adapted from ??*

### Establishing Drosophila RASopathy models

To generate a broad cross-section of RASopathy models, we stably expressed inducible transgenes for 13 RASopathy-associated human gene isoforms in flies: six *PTPN11* disease isoforms (Y279C, R498T, E510Q, D61G, N308D, E510P), three *RAF1* disease isoforms (S257L, L613V, D486G), two *BRAF* isoforms (T531, Q257R), and single isoforms of *KRAS* (G12D) and *HRAS* (G12S). Different isoforms are associated with different RASopathy, perhaps reflecting different signaling properties (Supplementary Figure 1). Using standard transgenic technology (Venken and Bellen, 2012), each transgene was inserted into the same genomic site to reduce expression differences due to insertion site. Each transgene was fused to a UAS-based inducible promoter that is silent until a GAL4 ‘driver’ is introduced. This system allows for both broad (*tub-GAL4*) and targeted expression including *ptc-GAL4* (directs expression in discrete regions of the developing wing, leg, and *eye imaginal* epithelia), *765-GAL4* (entire developing wing epithelia), and *byn-GAL4* (larval and adult hindgut). Higher ambient temperatures promote elevated GAL4 driver activity within the fly, a further step of controlling expression.

### RASopathy variants induced abnormal phenotypes

RASopathies can have both broad and more targeted or mosaic impact throughout the patient's body (Hafner and Groesser, 2013). We therefore explored the effect of expressing the RASopathy isoforms in restricted groups of cells as well as more broadly across the wing epithelia.

To explore overall lethality as a quantitative measure, we crossed each line to three different GAL4 drivers (*765-GAL4*, *ptc-GAL4*, and *byn-GAL4*) at four temperatures (22, 25, 27 and 29 °C; Figure 1B, Supplemental Figure 1). Survival was scored for pupariation and for adult eclosure, which provided a good framework for comparing phenotypes and also for the drug screens described below. Using a *ptc-GAL4* driver based on the *patched* (*ptc*) promoter, the transgenes were targeted to a discrete region of the developing wing epithelium (‘wing disc’). This discreet stripe within the developing epithelia gives rise to the adult wing tissue between longitudinal veins L3 and L4. Fly lines were examined at increasing temperatures to progressively elevate levels of *ptc-GAL4* activity (Figure 1B). Increased expression of *RAS*- and *RAF-*expressing lines at higher temperatures— 27 °C and 29 °C—led to lethality before adult stages. In contrast, the *PTPN11*-expressing lines survived to adulthood at these temperatures. Overall, the different transgenic lines showed significant variability with regard to lethality (Figure 1B, Supplemental Figure 1).

Focusing on specific disease isoforms, we found that *ptc-GAL4*-mediated expression of the activating mutant isoform RAF1^L613V^ (*ptc*>*RAF*^*L613V*^) led to a substantial increase in wing vein material within the *ptc* expression domain (inter-vein region between L3 and L4). This effect was further enhanced at 25 °C. Drosophila wing veins emerge at sites of high RAS pathway activity (Karim and Rubin, 1998), consistent with RAF1^L613V^ acting as an activated isoform. In contrast, *ptc*>*RAF1*^*D486G*^—containing a variant in RAF1’s DFG motif that controls kinase activity—led to mild but consistent loss of wing cross-veins within the *ptc* expression domain. This suggests that, in this context, *RAF1*^*D486G*^ acts as a loss-of-function allele. All *PTPN11* lines except one (*PTPN11*^*D61G*^) altered wing vein material patterns outside of the *ptc* expression domain (Figure 2A; Supplemental Figure 3). The strongest examples of this non-autonomous ectopic venation were observed in *ptc*>*PTPN11*^*R498W*^ wings (Figure 2A).

**Figure 2.**
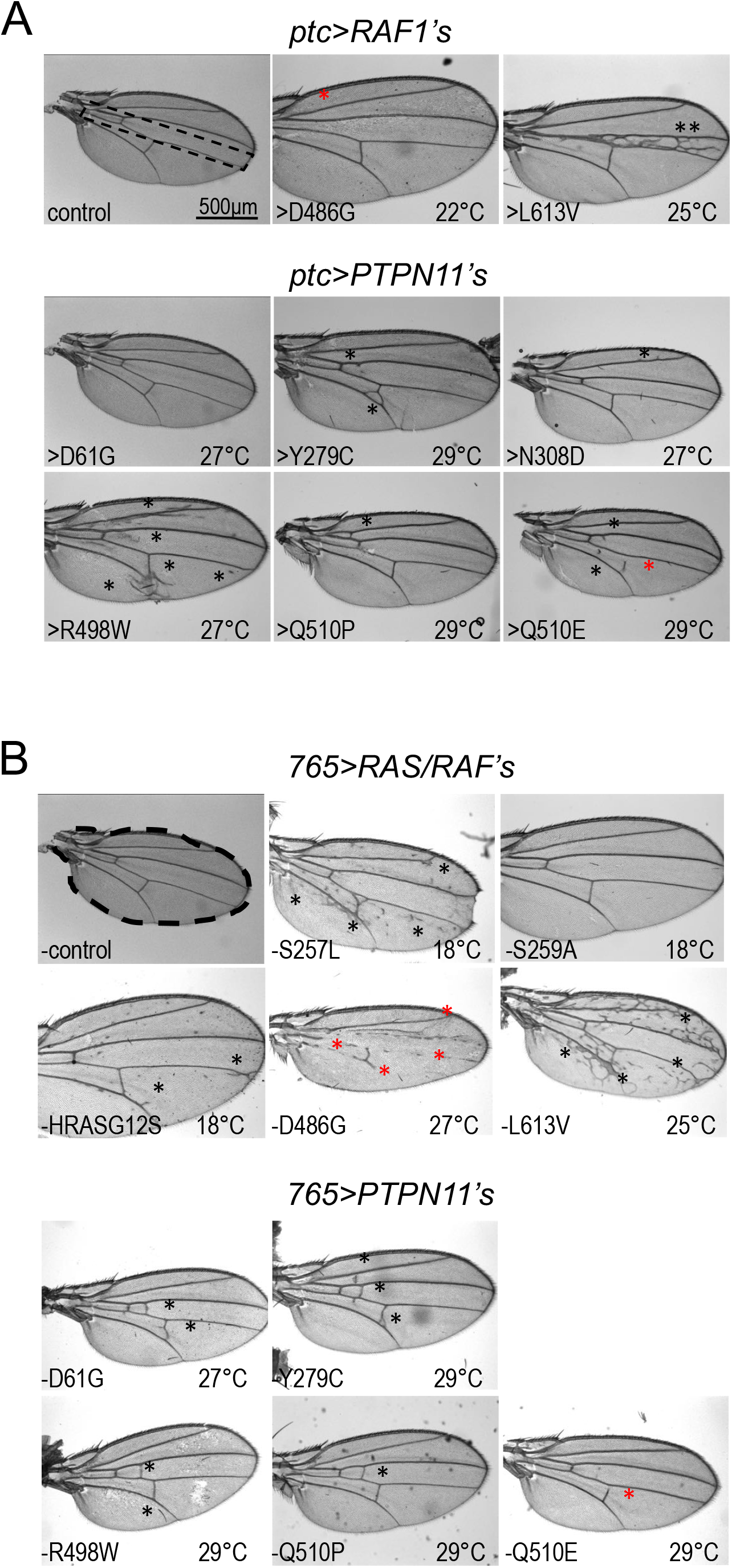
(A) Bright field images of adult fly wings in which RASopathy isoforms were overexpressed using the *ptc-GAL4* driver. Control wing with dotted outline indicates the region within which the *ptc-GAL4* driver is active. Upper panels: RAS/RAF isoforms-expressing flies exhibited ectopic wing venation within the *ptc* domain. Lower panels: *PTPN11* isoforms-expressing flies exhibited ectopic wing venation in different parts of the wing, but primarily outside the *ptc* domain. For panels in (A) and (B) the temperature at which the transgene was induced is indicated in each panel; the black asterisk indicates ectopic veins and the red asterisk indicates suppression of normal wing veins. Supplementary Figures 2-5 summarize these experiments showing the range of phenotypes at 18 °C, 22 °C, 25 °C, 27 °C, and 29 °C. (B) Bright field images of adult fly wings in which RASopathy isoforms were induced using *765-GAL4* driver. Control wing with dotted outline showing the region where the *765-GAL4* driver is active. Upper panels: RAS/RAF isoform-expressing flies mostly exhibited ectopic wing venation; the exception was *RAF1^D486G^*, which suppressed wing vein formation (red asterisks). Lower panels: *PTPN11* isoforms-expressing flies exhibited ectopic wing venation in different parts of the wing. *PTPN11* isoforms consistently induced a milder ectopic wing venation phenotype compared to the RAS/RAF isoforms.

We next explored the consequences of expressing the transgenes more uniformly across the larval wing epithelium using a *765-GAL4* driver (Figure 2B; Supplemental Figures 1B, 4, 5). Again, increased expression of RAS/RAF isoforms at higher temperatures—27 °C, 29 °C—led to lethality prior to eclosion as adults. *765*>*KRAS*^*G12D*^ and *765*>*BRAF*^*W531C*^ failed to survive even at lower temperatures, preventing analysis of their wing vein phenotypes. All *PTPN11* lines survived till adulthood. *765*>*RAF1*^*L613V*^ exhibited a strong temperature-dependent increase in wing venation; *765*>*RAF1*^*S257L*^ and *765*>*HRAS*^*G12S*^ lines exhibited a moderate increase in wing venation (25 °C).

Effect on wing veins were not restricted to a specific region of the wing. *765*>*RAF1*^*D486G*^, predicted to have loss of kinase activity, exhibited temperature-dependent suppression of wing veins in broad areas of the wing. By contrast, expression of the *PTPN11* transgenes with a *765-GAL4* driver led to a very specific effect: an increase in cross-veins between the longitudinal veins (L1-L4), with the penetrance of this phenotype being fairly low. In summary, expression of RAS or RAF disease isoforms led to ectopic wing veins throughout the wing field. Expressing *PTPN11* disease isoforms primarily drove ectopic wing cross-veins, suggesting a more cell type-specific effect of this class of RASopathy variants as has been reported in murine RASopathy models (Araki et al., 2004).

### RASopathy models responded differentially to therapeutics

As with many Mendelian diseases, RASopathy patients have limited therapeutic options. In our studies, we observed differences between the various RASopathy mutant isoforms with respect to their wing phenotypes. Here, we extend our studies to determine whether this heterogeneity of phenotypes extends to drug response. We screened a large panel of compounds that included FDA-approved drugs as well as polypharmacological tool compounds co-developed in our laboratory.

To screen for therapeutics, we used a quantitative Drosophila survival assay we previously developed to model various human cancer paradigms (Dar et al., 2012; Das et al., 2018; Das and Cagan, 2017). We used our lethality screen (Figure 1B, Supplemental Figures 1A) to identify 27 °C as the temperature that best yielded a minimal number of pupae or adults across all lines. This provided a large window for therapeutic rescue. We assessed a panel of 53 therapeutic compounds (Supplemental Figure 6) including clinically approved RAS/MAPK pathway inhibitors (*e.g.*, vemurafenib, axitinib), RTK inhibitors including inhibitors of RET (vandetanib, cabozantinib) and EGFR (erlotinib, lapatinib), SRC inhibitors, *etc*. We tested inhibitors targeting the proteasome, histone deacetylases (HDACs), and HSP90, which have shown promise as therapies in different RAS-dependent cancer paradigms. We also tested statins, which have been assessed in clinical trials (NCT02713945) on a small number of RASopathy patients.

For the drug screen, we used the driver *765-GAL4* at 27 °C. The ratio of the number of treated:untreated surviving pupae/adults (“% rescue”) was used to compare results across lines. Figure 3A provides a heat map of our drug screening results. The response of each model to therapeutics proved unique, with unexpectedly limited overlap of drug efficacy across different models (Figure 3A-C). The same panel of drugs were well-tolerated by control flies at the tested doses (Supplementary Figure 10).

**Figure 3.**
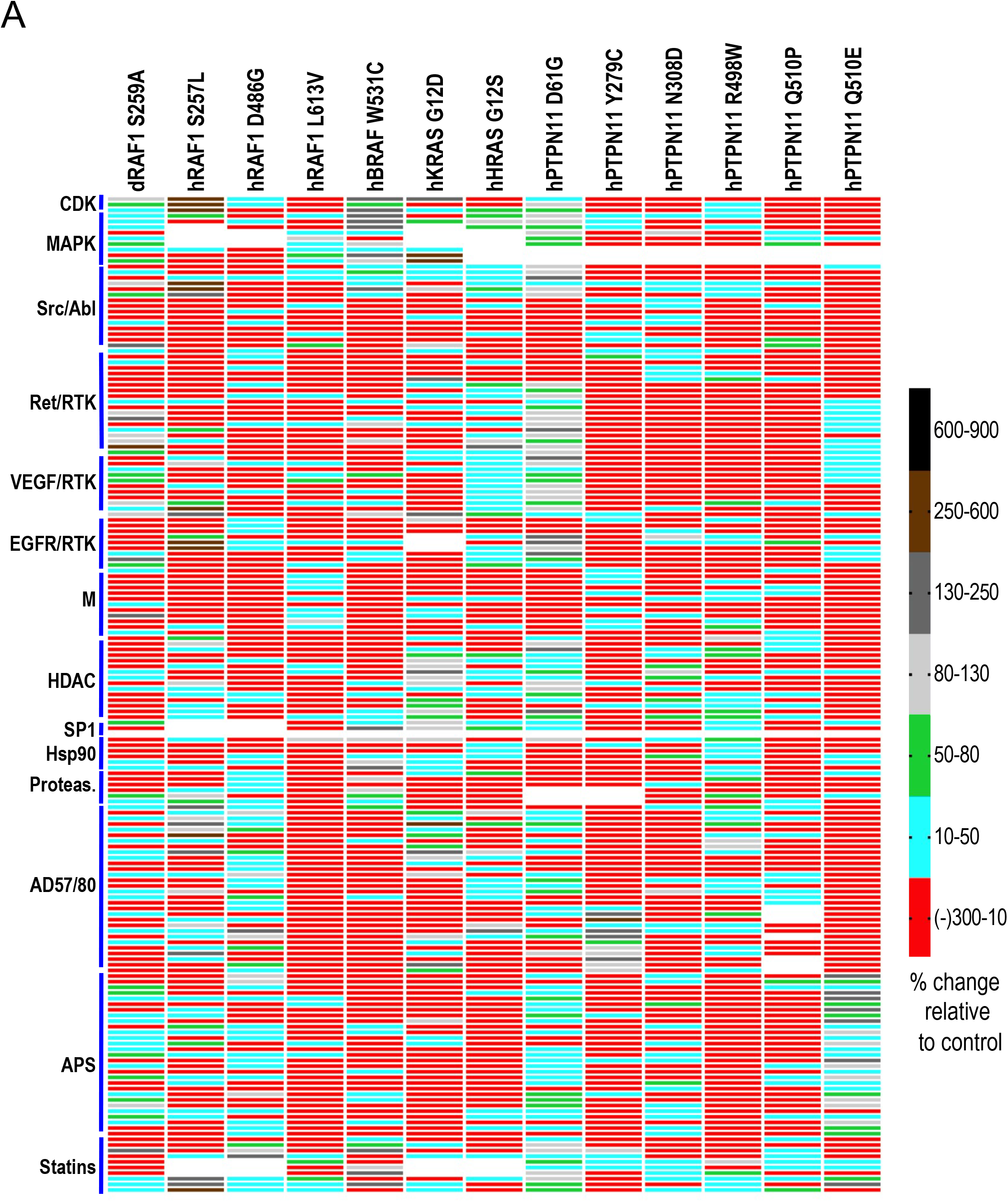
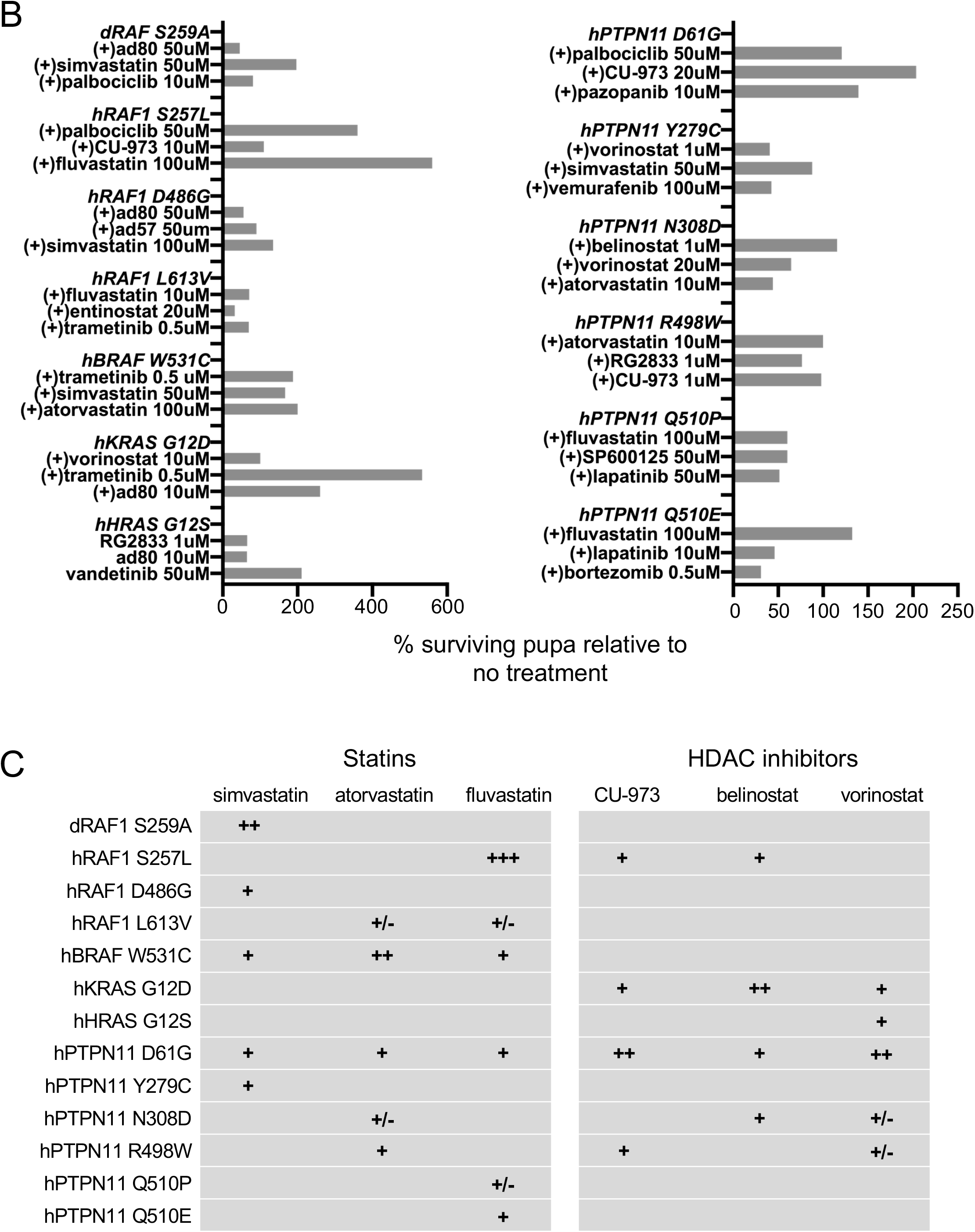
(A) Heat map depicting response of RASopathy models to a panel of indicated drugs and tool compounds. For each model, the heat map indicates the ratio of the number of pupae surviving following treatment compared to no treatment controls. This is represented as percent change compared to control as shown in the adjacent key. As in the previous figure, viability is assessed as the mean of four replicates for each condition. Each model exhibited a unique pattern of responses to the panel of drugs tested. The AD57/AD80 and APS family of tool compounds were developed in-house as previously published (Dar et al., 2012; Sonoshita et al., 2018). (B) Select top drug hits for each RASopathy model. Shown on the left are the RAS/RAF models and on the right the *PTPN11* models. As in (A), the bars represent the ratio of the number of pupae surviving following treatment compared to no treatment controls, represented as percentage change compared to control. No treatment controls often have slightly different survival rates (as indicated by error bars in Supplementary Figure 1A) and are therefore repeated for each batch of drugs tested (see methods), providing more accurate estimates of drug rescue in different experiments. (C) Table showing qualitative relative response of RASopathy model flies to statins and HDAC inhibitors. These two classes of compounds showed the broadest efficacy across the thirteen models tested. No single drug showed efficacy across all models. Statins showed better efficacy in RAS/RAF models compared to *PTPN11* models, while HDAC inhibitors showed the opposite.

***Statins*** affect cholesterol biosynthesis, which perturbs membrane localization of components of the RAS pathway. Three statins in our survey, atorvastatin, simvastatin, and fluvastatin, were active across the broadest palette of transgenic lines tested including members from both RAS/RAF and *PTPN11* class of models. At least one statin was active for each RASopathy model. Simvastatin and fluvastatin showed the broadest efficacy, improving viability of *RAF1*^*S259A*^, *RAF1*^*D486G*^, *BRAF*^*W531C*^, *PTPN11*^*D61G*^, *PTPN11*^*Y279C*^ flies (Figure 3A-C).

***Histone deacetylase inhibitors*** belong to a group of inhibitors that target various classes of HDACs (Chun, 2015; Duvic and Vu, 2007; Kaushik et al., 2015; Keller and Jung, 2014). HDAC inhibitors also showed fairly broad activity. The viability of *KRAS*^*G12D*^ flies were consistently improved by most HDAC inhibitors tested (vorinostat, entinostat, belinostat, RG2833). Viability of *PTPN11*^*D61G*^ flies were improved by vorinostat, entinostat, belinostat, and CUDC-9073 treatment. Viability of *RAF1*^*S257L*^ flies were improved by belinostat and CUDC-9073.

***AD and APS family*** of small molecule polypharmacological inhibitors target multiple kinases; they are especially potent RAS pathway inhibitors in cancer models (Dar et al., 2012; Sonoshita et al., 2018). These compounds were active against the *PTPN11*^*Q510E*^ but not the *PTPN11*^*Q510P*^ fly model, indicating a surprising specificity to a single amino acid change. We had previously described a polypharmacological class of drugs that targeted both cellular and lipid kinases (Dar et al., 2012). One member of that class, AD80, showed good activity and improved viability of multiple different models, including *RAF1*^*S257L*^ and *KRAS*^*G12D*^ strongly and *PTPN11*^*R498W*^ more modestly.

***CDK4/6 inhibitor*** palbociclib improved the viability of 5/7 RAS/RAF flies: *RAF1*^*S257L*^, *RAF1*^*S259A*^, *BRAF*^*W531C*^, *KRAS*^*G12D*^, and *HRAS*^*G12S*^ flies. Notably, palbociclib only increased viability of one *PTPN11* fly model, *PTPN11*^*D61G*^, indicating specificity for the RAS/RAF models.

***RAS/MAPK pathway inhibitors*** are another class of inhibitors that improved viability of some of the RASopathy models, primarily flies with the RAS/RAF variants. Trametinib improved viability of *KRAS*^*G12D*^ flies, vemurafenib rescued *BRAF*^*W531C*^ and *PTPN11*^*D61G*^ flies strongly and *RAF1*^*S257L*^, *RAF1*^*S259A*^ more modestly.

The differential response of the RASopathy fly models to different classes of drugs indicated that these models could be activating different patterns of signaling pathways *in vivo*. We next looked at the pattern of signaling for each RASopathy model in developing fly tissues, in the presence or absence of the top candidate drugs identified in our screen.

### Western blot analysis identified altered pathways

We recently published the use of an antibody panel to monitor multiple signaling pathways in both normal tissue and oncogenic models, including in the presence of therapeutics (Das et al., 2018; Das and Cagan, 2017). Here, we performed a similar, broader western blot analysis by gathering lysates from whole developing larvae where RASopathy variants had been transiently induced. We induced the RASopathy isoforms broadly in tissues of the developing Drosophila larvae to more closely mimic the germline nature of these variants and the observed pleiotropic effects in multiple tissue types (Figure 4A; *tubGAL4; GAL80^ts^>transgene*). Our panel focused on pathways with known roles in human disease. By comparing activity levels for each signaling pathway, we assessed how each RASopathy isoform activates distinct signaling pathways *in situ* (Figures 4, 5; Supplementary Figure 7).

**Figure 4.**
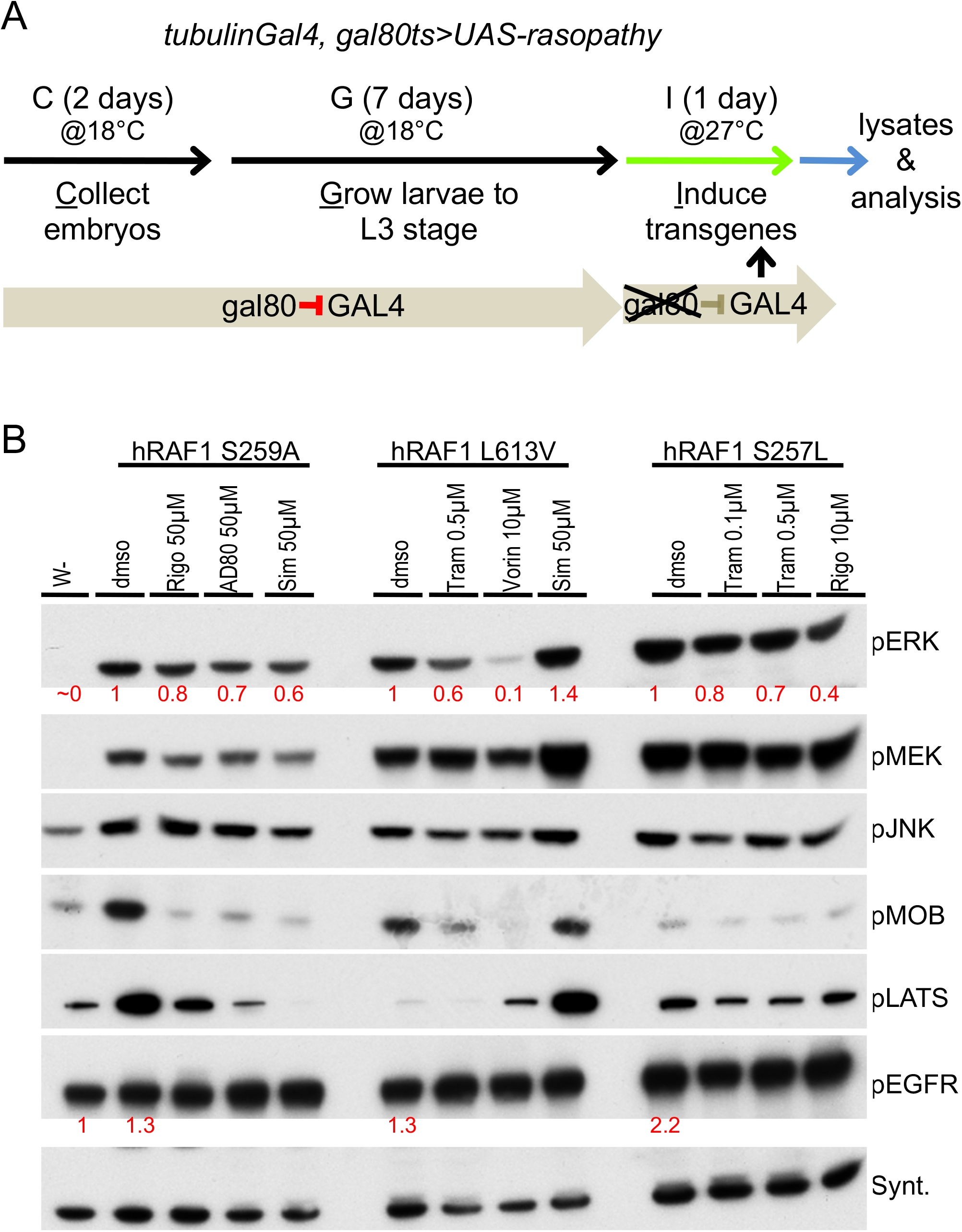
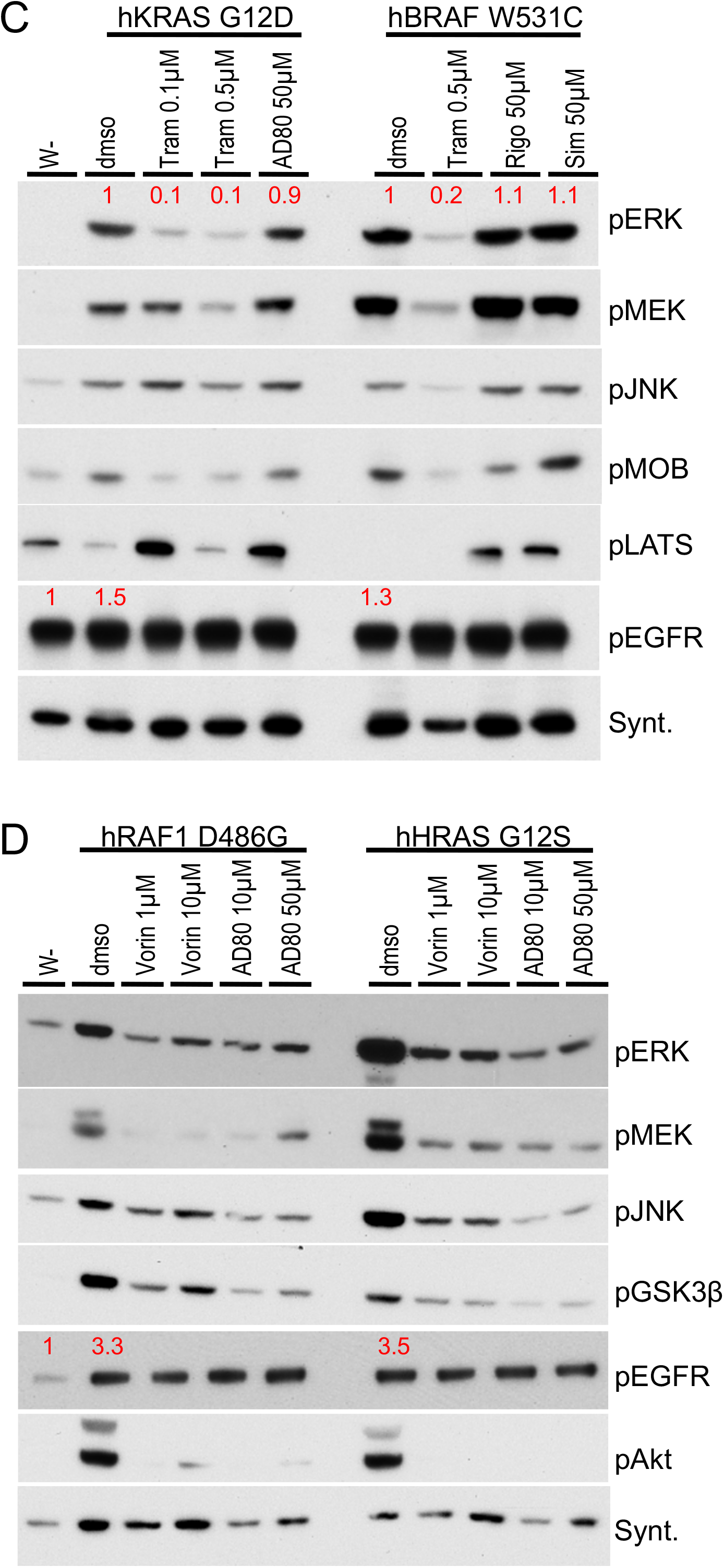
(A) Flowchart depicting timeline to induce expression of RASopathy isoforms in developing Drosophila larvae followed by western blot analysis. Embryos from flies were collected in a fixed timespan (Collection), and larva were allowed to develop at 18 °C until L3 stage (Growth). At this temperature transgene expression was not induced: basal expression of an included temperature sensitive GAL80-variant (GAL80^ts^) inhibited GAL4-dependent UAS-transgene activation. After reaching L3 stage the larvae were shifted to 27 °C, which led to destabilization of GAL80^ts^ protein and induced expression of the RASopathy encoding transgenes (Induction). After a fixed time of induction, larvae were collected and whole-body lysates extracted for western blot analysis. (B) Western blot analysis of indicated *RAF1* models. The first lane in this and subsequent panels represent lysates from *w-* control flies; DMSO represents treatment with the solvent in the absence of drug. Drug doses represent the condition at which the drugs showed efficacy in the screens in Figure 4. All *RAF1* models exhibited strong upregulation of pERK levels compared to control flies (lane 1; ~0); relative quantitation indicated below in red in this and subsequent panels. pJNK and pMEK levels were also increased by RAF1 isoforms (compare *w-* to DMSO lanes). Hippo pathway activity markers pMOB and pLATS were differentially regulated by the RAF1 isoforms; drug treatments led to clear effects on these markers in most *RAF1* lines tested. Downregulation of pMOBS and/or pLATS is predicted to promote cellular growth. (C) Western blot analysis of KRAS^G12D^ and BRAF^W531C^ isoforms. These isoforms induced strong upregulation of pERK and pMEK levels, moderate upregulation of pJNK levels and differential regulation of Hippo pathway markers pMOBS and pLATS (compare *w-* to dmso lanes). Treatment with MEK inhibitor trametinib suppressed pERK upregulation by both isoforms. Both isoforms suppressed the growth inhibitory Hippo pathway marker pLATS, while most drug treatments upregulated pLATS. (D) Western blot analysis of RAF1^D486G^ and HRAS^G12S^ isoforms, which induced strong upregulation of pERK, pMEK, pJNK, pAKT (PI3K pathway), and pGSK3β (Wnt/Wg pathway) levels (compare *w-* to dmso lanes). Vorinostat and polypharmacological compound AD80 suppress levels of all these markers. pERK, pMEK, pJNK, pAKT, pMOBS, pLATS, and pGSK3β indicate phosphorylated forms of the proteins. Syntaxin, in this and subsequent panels, was used as one method of assessing loading control; see Methods for full description.

#### RAS/MAPK pathway

We first assessed the level of activation of a key marker of the MAPK pathway, phosphorylated-ERK (pERK), to identify conditions in which most isoforms induce MAPK activity. All lines broadly expressing *RAS*/*RAF* isoforms exhibited a stable increase in levels of both pERK and pMEK (Figure 4). The *PTPN11* lines proved more complex. Most *PTPN11* lines displayed transient activation of pERK and basal levels of phosphorylated-MEK (pMEK; Figure 5, Supplementary Figure 7), consistent with previous work in mammalian models (Eminaga and Bennett, 2008; Zheng et al., 2018). Two lines—*PTPN11*^*Y279C*^, *PTPN11*^*Q510E*^—displayed higher levels of pMEK, indicating more complex regulation of MAPK activity in which pMEK and pERK do not mirror one another. This phenomenon has been reported in mammalian studies: recruitment of additional cofactors differentially altered phosphorylation levels of pERK *vs.* pMEK (Kuang et al., 2009; Lito et al., 2014; Ordan et al., 2018).

*PTPN11*^*Y279C*^, *PTPN11*^*Q510P*^ lines displayed initial activation of pERK followed by suppression of pERK within 24 hours, suggesting negative feedback regulation, a well-described feature of the MAPK pathway (Fritsche-Guenther et al., 2011; Lake et al., 2016; Shin et al., 2009). Still, activation of the pERK signal was not consistently observed, presumably due to the negative feedback regulation of the MAPK pathway (Figure 5; Supplementary Figure 7), which is another key point of difference with the *RAS*/*RAF* lines. The exception was *PTPN11*^*N308D*^, which activated pERK under all tested induction conditions with little or no observed negative feedback regulation of pERK, similar to the response of RAS/RAF isoforms (Figures 4, 5; Supplementary Figure 7).

In the course of these experiments we identified one condition in which all PTPN11 isoforms showed upregulation of pERK levels above baseline, and we used this condition for all subsequent analysis (see Methods).

#### EGFR activation

EGFR was moderately activated (pEGFR) by both the *RAS*/*RAF* (*RAF1*^*S257L*^, *RAF1*^*D486G*^, *HRAS*^*G12S*^) and *PTPN11* (*PTPN11*^*N308D*^, *PTPN11*^*Q510E*^, *PTPN11*^*Q510E*^) transgenes.

#### JNK/SAPK pathway

The *RAS*/*RAF* and *PTPN11* lines also displayed differences in their regulation of JNK pathway signaling. While 6/7 *RAS*/*RAF* lines (exempting *HRAS*^*G12S*^) strongly increased pJNK levels above basal levels, 4/6 *PTPN11* lines (exempting *PTPN11*^*N308D*^, *PTPN11*^*Q510E*^) decreased phosphorylated-JNK (pJNK) levels. Previous studies including our own with oncogenic RTK, RAS/MAPK, and SRC components identified increased JNK activity as a key aspect of transformation in Drosophila tissues; this study extends these observations (Das et al., 2013b; Das and Cagan, 2017; Rudrapatna et al., 2014; Vidal et al., 2006). Further, we provide evidence that multiple *PTPN11* RASopathy lines regulate the JNK pathway in a distinct manner.

#### Hippo pathway

Using the pathway activity markers pMOB (pMats) and pLATS (pWts), we found that *RAS*/*RAF* lines had a complex spectrum of Hippo pathway activation. *RAF1*^*S259A*^ increased levels of both markers. *RAF1*^*L613V*^, *KRAS*^*G12D*^, and *BRAF*^*W531C*^ increased pMOB but decreased pLATS. *RAF1*^*D486G*^ and *HRAS*^*G12S*^ displayed basal levels of both markers. Regarding *PTPN11*: *PTPN11*^*D61G*^ and *PTPN11*^*Y279C*^ showed decreased levels of only pLATS, *PTPN11*^*R498W*^, *PTPN11*^*Q510E*^, and *PTPN11*^*Q510P*^ showed decreased levels of pMOB. *PTN11*^*N308D*^ displayed decreased levels of both markers. These data suggests complex regulation across the Hippo pathway by different isoforms (see Discussion).

#### Other pathways

Two *RAS*/*RAF* lines, *RAF1*^*D486G*^ and *HRAS*^*G12S*^, demonstrated significantly increased levels of pAKT and pGSK3β, which were restrained by drugs that improved survival of these variants in our viability assays (Figure 4D). The PI3K-AKT signaling axis promotes growth, survival, invasion/metastasis and regulates energy homeostasis in vertebrate cells and in Drosophila (Dillon and Muller, 2010; Fruman et al., 2017; Hirabayashi et al., 2013; Witte et al., 2009). Deregulation of this pathway is associated with a variety of human diseases including cancer, diabetes, cardiovascular and neurological diseases. AKT signaling affects components of other signaling pathways including upregulation of phosphorylated GSK3β, which in turn leads to stabilization of β-catenin/Armadillo a key Wnt pathway transcription factor (Beurel et al., 2015; Das et al., 2013a; Hanahan and Weinberg, 2011; Valenta et al., 2012). Upregulation of these two interdependent markers is unique to *RAF1*^*D486G*^ and *HRAS*^*G12S*^, providing additional examples of how specific mutations can uniquely impact signaling networks that regulate cellular growth and homeostasis.

### Western blot analysis highlighted biomarkers of drug efficacy

Using a targeted screen of clinically relevant drugs and compounds, we identified a small set that improved viability of our RASopathy lines (*not shown*). Given the heterogeneity of pathway activity across different RASopathy lines, we used our western analysis to explore the activity of effective drug candidates for each RASopathy model. Drugs were analyzed at two concentrations for their effects on our panel of protein markers. The vehicle dimethylsulfoxide (DMSO) alone was used as a control. Our analysis identified examples of both expected and potentially novel mechanisms by which drugs promote viability of RASopathy variant expressing Drosophila larvae and, in this section, we highlight some of the notable examples.

#### Trametinib, AD80, and vorinostat efficacy correlated with reduced MAPK activation

The potent MEK inhibitor trametinib reduced levels of pERK protein in three of the RAS/RAF models (*RAF1*^*L613V*^, *KRAS*^*G12D*^, *BRAF^W531C^*); reduction of MAPK activation correlated with improved viability (Figure 4B-D). By contrast, *RAF1*^*S257L*^ larvae treated with trametinib showed only moderate suppression of MAPK components pERK and pMEK (Figure 4B), despite improved viability.

The polypharmacological drug AD80 inhibits the RAS/MAPK pathway through inhibition of multiple targets including RAF kinases (Dar et al., 2012). Lysates from *RAF1*^*D486G*^ larvae treated with AD80 showed a strong reduction of both pERK and pMEK levels. In contrast, *KRAS*^*G12D*^ larvae treated with AD80 showed moderate reduction of pERK and no effect on pMEK levels. This suggests that AD80 acts on these two lines through a somewhat different palette of targets.

Treatment with vorinostat resulted in strong suppression of pERK levels (*RAF1*^*L613V*^ and *HRAS*^*G12S*^ larvae) or moderate reduction of pERK and pMEK levels (*KRAS*^*G12D*^ and *RAF1*^*S259A*^ larvae). Vorinostat is an HDAC inhibitor with a broad palette of identified targets including members of the RAS/MAPK pathway (*e.g.* (Yu et al., 2003; Zhong et al., 2013)). Its suppression of MAPK activation in multiple RASopathy models and its strong record of efficacy in patients indicates that vorinostat’s ability to reduce RAS/MAPK signaling may prove useful therapeutically.

#### Statin efficacy correlated with reduced MAPK signaling and activation of the Hippo pathway

Inhibitors of the statin group of drugs improved viability in our screens for a large number of RASopathy models, suggesting potential broad utility. Regarding RAS/MAPK signaling, treatment with simvastatin reduced levels of pERK and pMEK in *RAF1*^*S259A*^ larvae and pERK levels in *RAF1*^*D486G*^ larvae. Interestingly a similar effect on the RAS/MAPK pathway was not observed in *RAF1*^*L613V*^ or *BRAF*^*W531C*^ larvae. Instead, these lines exhibited strong upregulation of pLATS levels and moderate activity of pMOB, two markers for activation of the Hippo signaling pathway (Figure 4B-D). Activation of Hippo pathway signaling is known to have several effects, most notably reduced cell proliferation and tissue size (Davis and Tapon, 2019; Udan et al., 2003; Wu et al., 2003).

#### Drug efficacy in PTPN11 models correlated with growth-inhibitory activity of the Hippo pathway

Our western blot analysis above indicated that the *PTPN11* lines promoted only transient activation of the RAS/MAPK pathway, but more consistent regulation of the SAPK/JNK and Hippo pathways. We, therefore, monitored the latter two pathways in *PTPN11* larvae after treatment with drugs that increased viability (Figure 5A-B).

**Figure 5.**
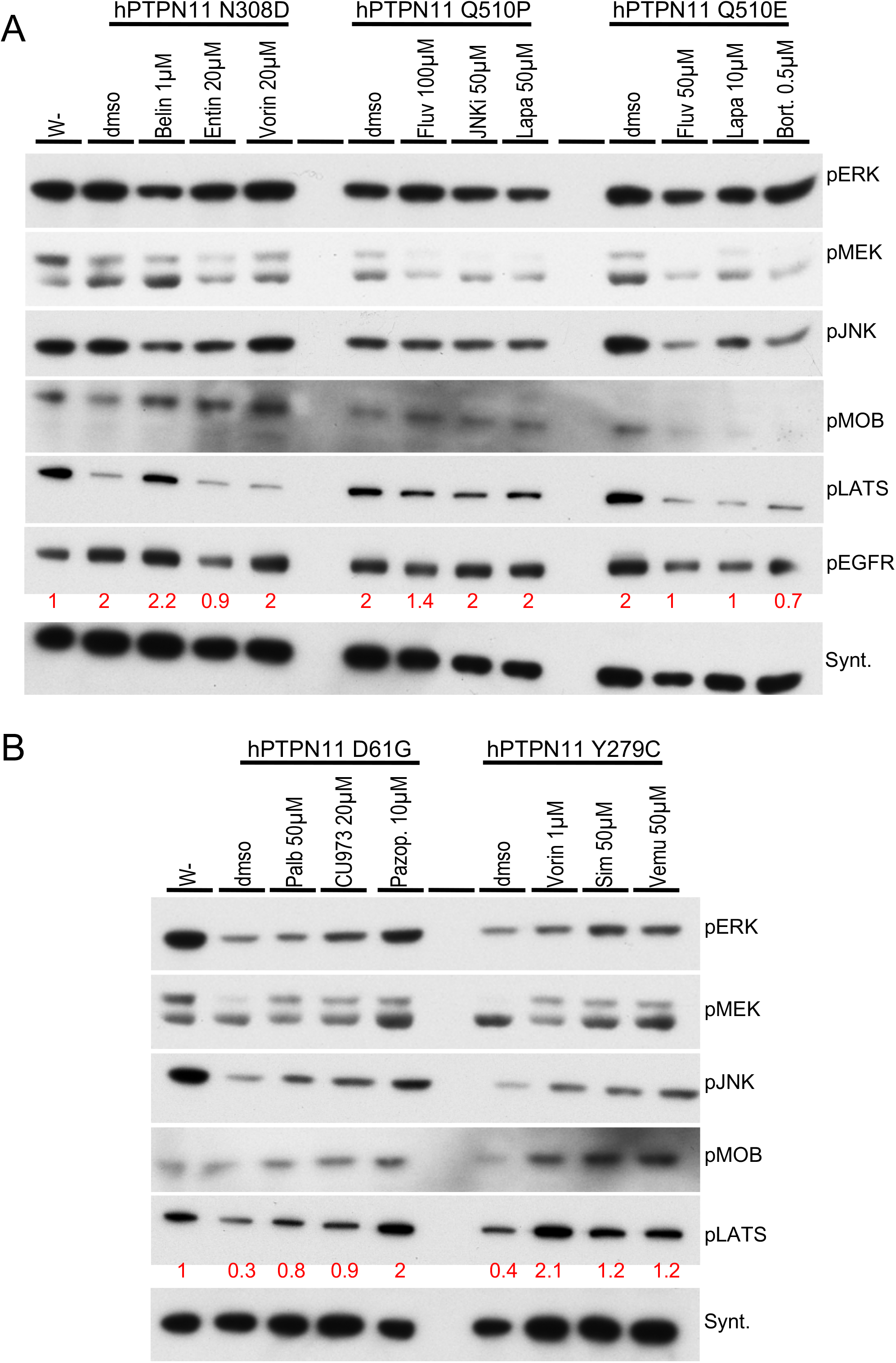
(A) Western blot analysis of indicated *PTPN11* models, which had a mild effect on the MAPK pathway (also see Supplementary Figure 7). All three isoforms induced almost two-fold upregulation of pEGFR levels; some drug treatments suppressed this induction. These isoforms showed differential regulation of growth inhibitory Hippo pathway marker pLATS and pMOBS. Notably, *PTPN11*^*N308D*^ lines exhibited reduced levels of these markers while drug treatments reversed that effect and upregulated one or both markers. (B) Western blot analysis of indicated *PTPN11* models. These two isoforms consistently reduced levels of growth inhibitory Hippo pathway markers pLATS and pMOBS; drug treatments reversed that effect, leading to upregulation of one or both markers.

For drugs effective at increasing viability of *PTPN11*^*D61G*^ (palbociclib, CUDC-973, pazopanib), *PTPN11*^*Y279C*^ (vorinostat, simvastatin, vemurafenib), *PTPN11*^*N308D*^ (belinostat, entinostat, vorinostat), or *PTPN11*^*Q510P*^ (fluvastatin, SP610025, lapatinib) lines, efficacy correlated with upregulation of growth-inhibitory markers of the Hippo pathway, pLATS and pMOBS (Figure 5A-B). For drugs that increased viability of *PTPN11*^*Q510E*^ lines (fluvastatin, lapatinib, bortezomib), efficacy correlated with suppression of pJNK levels. These results are consistent with our pathway analyses highlighting a potential role for Hippo and SAPK/JNK activity in RASopathy defects. Linking our observation that pEGFR activity was increased in a subset of *PTPN11* models—*PTPN11^N308D^*, *PTPN11^Q510P^*, *PTPN11*^*Q510E*^—we found that some successful therapeutics in these models restrained pEGFR activation (Figure 5A-B) including EGFR-inhibitors like lapatinib (for *PTPN11*^*Q510E*^). In summary, drugs effective in the *PTPN11* models represent a wide range of inhibitor classes, suggesting that EGFR, SAPK/JNK and Hippo pathway markers serve as common surrogate markers of treatment efficacy.

### Reducing HDAC1 in KRAS and RAF1 models phenocopied vorinostat treatment

In summary, our analyses identify key pathway biomarkers that could be useful to monitor treatment efficacy in future studies. They emphasize, however, the challenge of identifying a single therapeutic that is effective against most RASopathy-associated variants. A couple of notable exceptions were HDAC inhibitor vorinostat as well as statins, which were effective at improving viability of both RAS/RAF and *PTPN11* fly lines. Both classes of drugs have a wide range of clinical use in multiple disease paradigms, and the statins in particular are well tolerated over long periods of treatment.

To confirm that vorinostat’s rescue activity was linked to HDAC inhibition, we reduced activity of the Drosophila HDAC1 ortholog Rpd3 in lines expressing KRAS^G12D^ and RAF1^L613V^ throughout the developing wing. Expression of RAF1^L613V^ isoform displayed a temperature-dependent strengthening of the ectopic wing venation phenotype (Figure 2; Supplementary Figures 1, 2, 4). At 20 °C, targeted RNA interference-mediated knockdown of Rpd3 in *RAF1*^*L613V*^ lines (*765*>*RAF1*^*L613V*^, *rpd3-RNAi*) resulted in an increase of flies eclosing as adults (increase from 5% to 20%), as well as suppression of ectopic wing venation pattern (Figure 6A; also see Figure 2 and Supplementary Figure 4). At 25 °C, knockdown of Rpd3 did not suppress ectopic wing venation, but there was a consistent enlargement of overall wing blade size in *765*>*RAF1^L613V^, rpd3-RNAi* adults compared to *765*>*RAF1*^*L613V*^ flies (Supplementary Figure 8A). Suppression of these different phenotypic effects by knockdown of Rpd3 establishes the histone deacetylase pathway as a regulator of RAF1^L613V^-mediated signaling.

Furthermore at 20-25 °C, none of the *765*>*KRAS*^*G12D*^ flies survived to adult stages (Figure 2; Supplementary Figures 1, 4, 5). At 20 °C, knockdown of Rpd3 improved survival and resulted in 25% viable adult flies eclosing with normal wing veins (Figure 6B; Supplementary Figure 8B). At 25 °C, knockdown of Rpd3 (*765*>*KRAS^G12D^, rpd3-RNAi*) also improved survival and led to 10% of *765*>*KRAS*^*G12D*^ animals surviving to adulthood (Figure 6B). The wings of eclosed *765*>*KRAS*^*G12D*^, *rpd3-RNAi* adults, at 25 °C, still displayed excess venation similar to other RAS/RAF models indicating that Drosophila HDAC1 did not fully regulate all aspects of RAS pathway activity leading to vein formation. Taken together with our drug screen, it indicates that in *KRAS*^*G12D*^-expressing models vorinostat is more effective on pathways that regulate survival vs. wing venation. In summary, these genetic knockdown experiments indicated that HDAC proteins normally function to promote KRAS^G12D^ and RAF1^L613V^ activity in Drosophila.

**Figure 6.**
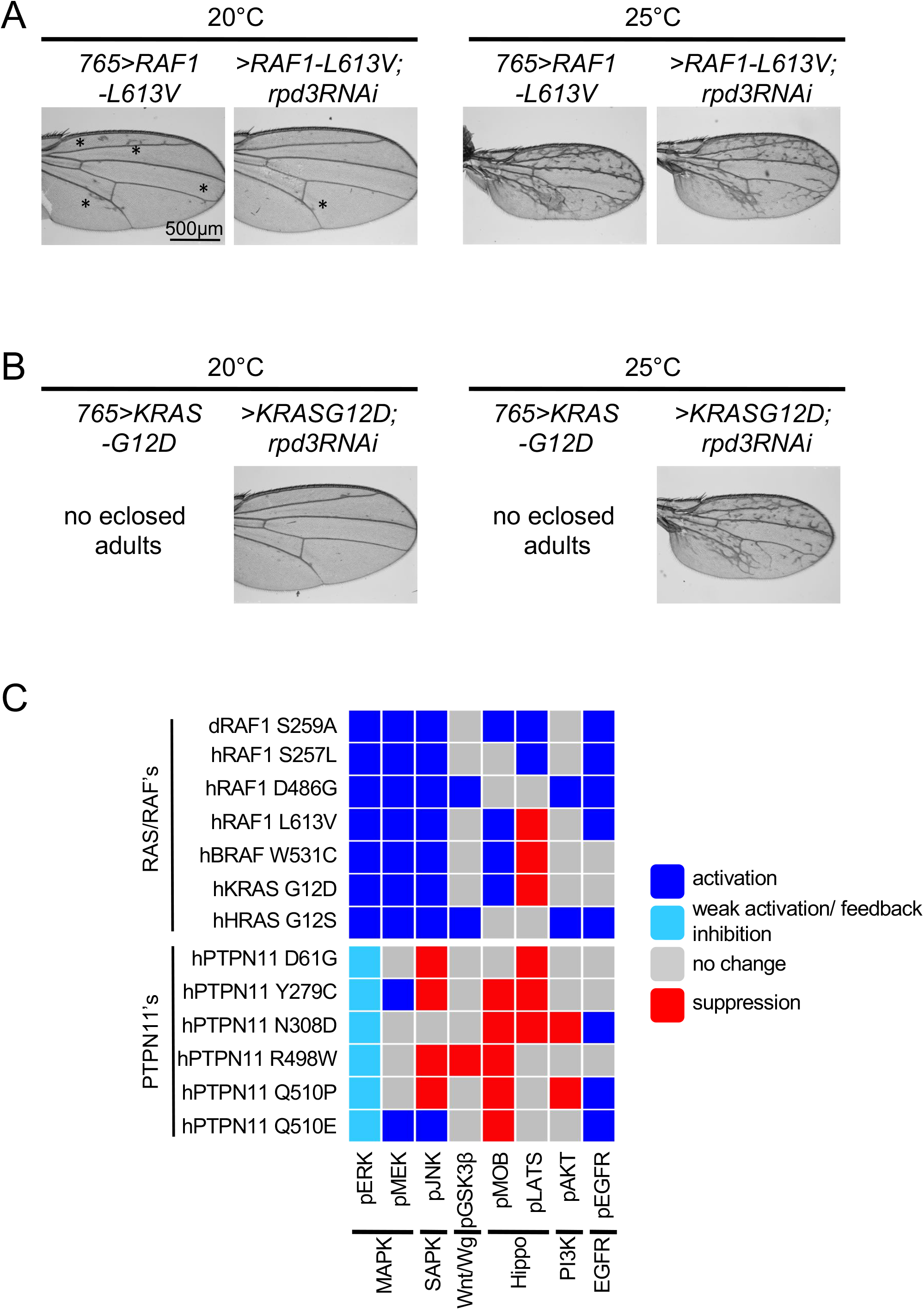
(A) Genetic modifier experiments with lines expressing *RAF1*^*L613V*^ demonstrated dependency on HDAC1. *RAF1*^*L613V*^ was expressed throughout the developing larval wing disc using the *765-GAL4* driver under different temperature conditions (also see Figure 2 and Supplementary Figure 2). In *765>RAF1*^*L613V*^ flies, ectopic wing venation phenotypes were observed at 20 °C, which was suppressed by RNAi-mediated knockdown of fly HDAC1 ortholog Rpd3 (rpd3-RNAi). The suppression of ectopic wing venation did not occur at 25 °C with stronger induction of the isoform. Black asterisk indicates ectopic veins. (B) Genetic modifier experiments with KRAS^G12D^ isoform demonstrate dependency on HDAC1 activity. When *765*>*KRAS*^*G12D*^ flies were raised at 20 °C and 25 °C, no adults eclosed. This developmental lethality was suppressed by co-expression of *rpd3-RNAi*; of note, the number of UAS transgenes is increased by one, which could affect expression levels. At 20 °C *765*>*KRAS*^*G12D*^, *rpd3-RNAi* flies exhibited near-normal wing vein pattern, while at 25 °C the ectopic wing venation pattern was not suppressed, presumably due to stronger induction of the isoform. (C) Summary of the pathway activation/signaling analysis of the different RASopathy models. Overall, RAS/RAF isoforms were significantly distinct from *PTPN11* isoforms in their activation of the MAPK pathway. However more broadly, each RASopathy isoform displayed a unique profile of regulation of major cellular pathways as assessed by the indicated markers.

## Discussion

Since the first clinical identification of RASopathy patients (Noonan and Nadas, 1958) a large effort has gone towards further defining and expanding the repertoire of genetic mutations associated with this syndrome. Identifying the genes responsible for Noonan syndrome (Tartaglia et al., 2001) and establishing that these genetic variants were known from their impact on cancer (Tartaglia and Gelb, 2010, 2005) were key to begin exploring the molecular mechanisms underlying the disease. Development of RASopathy animal models has provided important functional insights in the etiology of craniofacial, cardiac, and other developmental abnormalities associated with the disease (Hernández-Porras and Guerra, 2017; Jindal et al., 2015; Patterson and Burdine, 2020). These models have provided important molecular insights, for example that both gain-of-function and loss-of-function of different genes could lead to similar phenotypic outcomes (Bonetti et al., 2014; Patterson and Burdine, 2020; Stewart et al., 2010). Taken together, animal models have contributed immensely to our understanding of specific genetic variants and the underlying molecular mechanisms associated with RASopathy disease progression (Jindal et al., 2015).

Here, we provide a complement to these earlier studies: a systematic, side-by-side comparison of 13 RASopathy variants in a whole animal setting to explore similarities as and differences in biology, signaling, and response to candidate therapeutics. Expressing human RASopathy isoforms of *KRAS, HRAS, RAF1, BRAF*, and *PTPN11* in Drosophila, we report important differences between disease isoforms including distinct signaling pathways and drug responses (Figure 4C, Figure 7C). The differential regulation of pMOB vs pLATS in the various lines provides an instructive example. LATS is the key kinase that regulates phosphorylation and subsequent nuclear translocation of the downstream nuclear factor YAP/TAZ, a process that keeps cellular growth in check. MOB is an adaptor protein that regulates assembly of two branches of the Hippo pathway: the LATS complex and the NDR1/2 complex (Bichsel et al., 2004; Hergovich et al., 2006, 2005). Thus, pMOB levels are a dynamic readout of the activity of these two branches of the Hippo pathway (Kulaberoglu et al., 2017). Reduced levels of pLATS, a core component of the MST-LATS signaling cascade, is predicted to promote cellular growth, a pro-tumorigenic outcome. Indeed, genetic screens in vertebrate cells have shown that the RTK/RAS/MAPK pathway can act directly on LATS and YAP/TAZ phosphorylation status (Azad et al., 2020); increased levels of pMOB would indicate an activation of NDR1/2 pathway independent of the pLATS branch. This mechanism may explain the differential regulation of pMOB *vs*. pLATS that has emerged in our studies.

Matching drug response to disease isoform was a key goal of our work. For example, we demonstrated differences in steady-state RAS pathway signaling between *PTPN11* models and RAS/RAF models; these differences presumably underpin differences in drug response. Despite these differences, we identified HDAC inhibitors and statins as demonstrating efficacy across multiple disease isoforms including *KRAS, HRAS, RAF1,* BRAF, and *PTPN11*. Our genetic studies indicate that, at least for HDAC inhibitors, this suppression of whole-body defects is on-target. Together, these data emphasize the complexity between RASopathy isoforms but also provides a candidate roadmap to handle this complexity across disease subtypes. This, in turn, would allow more inclusive patient recruitment, an important advantage in rare Mendelian diseases.

Developing therapies for rare Mendelian diseases presents several challenges. Developmental abnormalities resulting from a RASopathy may not be fully reversed or rectified through therapy. Nevertheless, an important goal can be to broadly suppress aberrant signaling in the patient’s tissues, which in turn may provide measurable clinical and quality of life benefits. Chronic treatment of patients, especially young children, has whole-body risks including further damage to developing systems. For early-onset diseases such as RASopathies, chronic therapies are required that slow disease progression while protecting normal postnatal development. We address this challenge by emphasizing whole-animal screening of our Drosophila RASopathy models. Whole-animal screening often identifies hits that are different than those found in *in vitro*, *in silico*, or in cell line assays. HDAC inhibitors provide a useful example: they have broad effects across cellular networks, but reduced RASopathy-associated phenotypes across most of our RASopathy models. The result was improved viability and reduced RAS pathway activity.

From a practical standpoint, the small number of RASopathy patients that have a specific gene variant makes recruitment for clinical trials difficult. Although one of the more common Mendelian diseases, RASopathies as a class have proven difficult to execute clinical trials in part for this reason. One solution is to identify a set of biomarkers for assessing efficacy of candidate treatments as well as disease progression; our platform matches mutations to phenotypic severity and to drug response. A second approach is to identify therapies that are effective across a broad palette of RASopathy variants, a key goal of this study. Our data points to HDAC inhibitors and statins as candidates to fulfill this criterion: as a class of drugs each showed efficacy across most of our models. In preclinical studies, HDAC inhibitors have shown initial promise in suppressing hypertrophic cardiomyopathy, a common source of morbidity and mortality in RASopathy patients (Ferguson et al., 2013). HDAC inhibitors such as vorinostat and belinostat have previously been used in pre-clinical models to reduce RAS pathway signaling by multiple laboratories including our own, most commonly as part of a cancer drug combination (Das et al., 2018; He et al., 2019; Malone et al., 2017; Morelli et al., 2012; Yamada et al., 2018).

Statins collectively refer to a family of drugs that inhibit a key enzyme of the lipid biosynthesis pathway, HMG-CoA reductase, and are commonly used to lower the risk of heart failure by preventing myocardial infarction. In addition, pre-clinical data has demonstrated that statins reduce prenylation, a required step in localization and activity of RAS proteins. Statins have therefore been explored in various human disease paradigms that involve deregulation of RAS/MAPK and related signaling axes (Davies et al., 2016; Yang et al., 2020; Youssef et al., 2002). Importantly, we found that, for some RASopathy variants, the efficacy of some drugs including statins correlated with suppression of pathways outside the canonical RAS pathway. Examples include the Hippo and JNK pathways. These observations open the possibility that signaling networks outside the canonical RAS pathway may prove useful as therapeutic targets in specific disease variants. This mirrors work in a preclinical *PTPN11*^*D61Y*^ mouse model, which showed similar activation of pathways outside of RAS including PI3K/AKT/mTOR and JAK/STAT signaling (Altmüller et al., 2017). More recently, work on a rare RASopathy variant in the *RRAS2* gene reported dysregulation of the Hippo pathway (Capri et al., 2019; Nussinov et al., 2018). Future studies can dissect whether other RASopathy variants can regulate other non-RAS/MAPK pathway networks.

In summary, our results demonstrate important differences in signaling between different RASopathy-associated variants. Each RASopathy variant showed important differences in cellular signaling pathways, presumably accounting for unique phenotypic signatures including drug response. These differences mirror the broad range of morbidities presented by RASopathy patients. Despite these differences, our data highlights HDAC inhibitors and statins as having the ability in at least one pre-clinical model to have broader therapeutic impact. These classes of drugs have strong clinical histories including for extended use, making them interesting candidates for further exploration.

## Acknowledgments

We thank the Cagan Laboratory for important discussions.

## Funding

This work was supported by NIH-U54OD020353, NIH-R35HL135742.

## Competing interests

BG declares royalties from GeneDx, Correlegan. LabCop and Prevention Genetics.

## Data and materials availability

All data and accession numbers needed to evaluate the conclusions in the paper are present in the paper and/or the Supplementary Materials. Additional data related to this paper may be requested from the authors.

## Methods

### Antibodies

Antibodies used for Drosophila western blot analysis were: anti-pJNK; anti-pAKT; anti-pMOB; anti-pEGFR; anti-pMEK; anti-pLATS; anti-pGSK3β - (Cell Signaling); anti-pERK - (Sigma); anti-Actin; anti-Syntaxin; anti-ß-tubulin - (Developmental Studies Hybridoma Bank). Except anti-Syntaxin, all other antibodies were developed against human protein and various previous studies, by our group as well as others, have shown they cross react and can identify the Drosophila protein.

### Comprehensive Statistical Analysis

For pupal and adult viability analysis in Figure 1, mean and standard error of the mean (SEM) were calculated and 4-5 vials/experiment (biological replicates) per condition were analyzed. Each vial contained between 20-80 developing embryos. For the large ~60 drug library screen in Figure 4, four vials per drug (biological replicates) with approximately 20–40 embryos per vial were analyzed by aliquoting a slurry of collected embryos in embryo compatible buffer (Das et al., 2013a; Na et al., 2013). For this large screen, absolute numbers of surviving pupae/adults were compared to no drug treatment to obtain a ratio of drug treatment/no drug treatment (percentage increase over baseline). To assess statistical significance of difference between means, t-Test with Welch’s correction was performed using PRISM software. The correction was used to account for samples with unequal variances and unequal sample sizes. Candidate drugs showing the highest efficacy (drug conditions with *p* values ranging from 0.05 to 0.3) in this primary screen were re-tested in a secondary test in a similar manner, except in addition total number of starting embryos in each vial was visually counted. This allowed for precise quantitation of increased viability by comparing the proportion of pupa/embryos rescued, with or without drug. Statistical significance, p<0.05, of difference between means, t-test with Welch’s correction was performed using PRISM software in these retests.

### Generating Drosophila Transgenic RASopathy Models

The cDNAs corresponding to RAS/MAPK pathway gene variants found in patients were subcloned into Drosophila transformation vector pUAST-attB. cDNA’s were obtained from commercially available (AddGene) sources and variants were introduced by performing overlapping PCR. Internal primers with altered DNA sequence were combined with primers at the start and end of the corresponding cDNA which contained restriction enzyme overhangs. Injection and creation of attp40 transgenics was done by BESTGENE Inc. Flies corresponding to *UAS-Rpd3* knockdown lines (TRIP lines) were obtained from Bloomington Drosophila Stock Center. The following primers were used:

**hRAF1(S257L)**
F-hRAF1-kzk-EcoR1-Start
GAATTCCAAAACATGGAGCACATACAGGGAGCTTGG
F-hRAF1-767(S257L)
GGTTGACATCCACACCTAATGTCCAC
R-hRAF1-756(S257L)
CATTAGGTGTGGATGTCAACCTCTGCCTCTGG
R-hRAF1-XbaI-Stop
TCTAGACTAGAAGACAGGCAGCCTCGGGG

**hRAF1(L613V)**
F-hRAF1-kzk-EcoR1-Start
GAATTCCAAAACATGGAGCACATACAGGGAGCTTGG
F-hRAF1-1835(L613V)
CTGTACCGAAGATCAACCGGAGCGC
R-hRAF1-1822(L613V)
GTTGATCTTCGGTACAGAGTGTTGGAGCAG
R-hRAF1-XbaI-Stop
TCTAGACTAGAAGACAGGCAGCCTCGGGG

**hRAF1(D486G)**
F-hRAF1-kzk-EcoR1-Start
GAATTCCAAAACATGGAGCACATACAGGGAGCTTGG
F-hRAF1-1454(D486G)
GAGGTTTTGGTTTGGCAACAGTAAAGTC
R-hRAF1-1437(D486G)
TGCCAAACCAAAACCTCCAATTTTCACTGTTAAG
R-hRAF1-XbaI-Stop
TCTAGACTAGAAGACAGGCAGCCTCGGGG

**hBRAF(W531C)**
F-hBRAF-kzk-NotI-Start
GCGGCCGCCAAAACATGGCGGCGCTGAG
F-hBRAF-1651(W531C)
GTGTTGTGAGGGCTCCAGCTTGTATC
R-hBRAF-1637(W531C)
GGAGCCCTCACAACACTGGGTAACAATAGC
R-hBRAF-XbaI-Stop
TCTAGATCAGTGGACAGGAAACGCACCATATCC

**hKRAS(G12D)**
F-hKRAS-kzk-NotI-Start
GCGGCCGCCAAAACATGACTGAATATAAACTTGTGGTAGTTGGAGCTGATGGCG
R-hKRAS-XhoI-Stop
GAGCTCTTACATAATTACACACTTTGTCTTTGAC

**hHRAS(G12S)**
F-hHRAS-kzk-NotI-Start
GCGGCCGCCAAAACATGACGGAATATAAGCTGGTGGTGGTGGGCGCCTCCGGTGT
R-hHRAS-XbaI-Stop
TCTAGATCAGGAGAGCACACACTTGCAGCTCATGCAGCCGGG

**hPTPN11(D61G)**
F-hPTPN11-kzk-EcoR1-Start
GAATTCCAAAACATGACATCGCGGAGATGGTTTC
F-hPTPN11-179(D61G)
GTGGTTACTATGACCTGTATGGAGGG
F-hPTPN11-164(D61G)
CATACAGGTCATAGTAACCACCAGTGTTCTGAATC
R-hTPTN11-XbaI-Stop
TCTAGATCACAGATCCTCTTCAGAGATGAGTTTTCTG

**hPTPN11(Y279C)**
F-hPTPN11-kzk-EcoR1-Start
GAATTCCAAAACATGACATCGCGGAGATGGTTTC
F-hPTPN11-833(Y279C)
GATGTAAAAACATCCTGCCCTTTGAT
R-hTPTN11-825(Y279C)
CAAAGGGCAGGATGTTTTTACATCTATTTTTG
R-hTPTN11-XbaI-Stop
TCTAGATCACAGATCCTCTTCAGAGATGAGTTTTCTG

**hPTPN11(N308D)**
F-hPTPN11-kzk-EcoR1-Start
GAATTCCAAAACATGACATCGCGGAGATGGTTTC
F-hPTPN11-919(N308D)
GCAGATATCATCATGCCTGAATTTGAAAC
R-hPTPN11-907(N308D)

R-hTPTN11-XbaI-Stop
TCTAGATCACAGATCCTCTTCAGAGATGAGTTTTCTG

**hPTPN11(R498W)**
F-hPTPN11-kzk-EcoR1-Start
GAATTCCAAAACATGACATCGCGGAGATGGTTTC
F-hPTPN11-1498(R498W)
GTGTGGTCTCAGAGGTCAGGGATG
R-hTPTN11-1479(R498W)
ACCTCTGAGACCACACCATCTGGATG

R-hTPTN11-XbaI-Stop
TCTAGATCACAGATCCTCTTCAGAGATGAGTTTTCTG

**hPTPN11(Q510E)**
F-hPTPN11-kzk-EcoR1-Start
GAATTCCAAAACATGACATCGCGGAGATGGTTTC
F-hPTPN11-1525(Q510E)
GCAGAGTACCGATTTATCTATATGGCG
R-hTPTN11-1513(Q510E)
GATAAATCGGTACTCTGCTTCTGTCTGGAC
R-hTPTN11-XbaI-Stop
TCTAGATCACAGATCCTCTTCAGAGATGAGTTTTCTG

**hPTPN11(Q510P)**
F-hPTPN11-kzk-EcoR1-Start
GAATTCCAAAACATGACATCGCGGAGATGGTTTC
F-hPTPN11-1525(Q510P)
GCACCGTACCGATTTATCTATATGGC
R-hTPTN11-1513(Q510P)
GATAAATCGGTACGGTGCTTCTGTCTGGAC
R-hTPTN11-XbaI-Stop
TCTAGATCACAGATCCTCTTCAGAGATGAGTTTTCTG

### Inhibitor Studies in Flies

Drugs were obtained from LC laboratories or Selleck Chemicals and were dissolved in dmso as stock solutions ranging from 1-200mM. Drugs (500–1000 μl) were diluted in molten (~50–60 °C) enriched fly food, aliquoted into 5-ml vials to obtain the final drug concentration in food. Based on previous analysis, after consumption of drug-containing food by larvae the circulating concentration of drug is 100–1000 fold lower than the concentration in fly food (Bangi et al., 2016; Das et al., 2018). 30–60 embryos of each genotype were raised on drug-containing food until they matured as third-instar larvae (whole larvae for western blot assay) or allowed to proceed to adulthood (viability assay and wing vein quantitation assay).

### Western Blot of Whole Larval Lysates and Quantitation

Three third-instar larva of each genotype (*tubulin-GAL4*; *gal80*^*ts*^ > *UAS-transgene*) were dissolved in Lysis Buffer (50 mM Tris, 150 mM NaCl, 1% Triton-X100, 1 mM EDTA) supplemented with protease inhibitor cocktail (Sigma) and phosphatase inhibitor cocktail (Sigma). Total protein in each sample was quantitated using BIORAD protein assay. Total protein amounts in each lysate was established by performing Bradford assay (BIORAD), and equivalent amounts (2–10 μg) of total protein was loaded per lane. As many of the isoform activated pathways are known to act on housekeeping proteins such as Syntaxin, Actin, and Tubulin, we relied on initial protein quantitation for accurate loading (Das et al., 2018, 2013a, 2013b; Das and Cagan, 2017). During western blot development we assessed, when possible, all three markers mentioned above. As expected, we found that different loading controls (Syntaxin *vs*. tubulin) were regulated differentially by RASopathy variant activation, and therefore initial protein quantitation (Bradford/BIORAD) is the more reliable method of ensuring accurate loading (Supplementary Figure 9). Samples were resolved on Invitrogen NU-PAGE gradient SDS-page and transferred by standard protocols. Membranes were stripped with SIGMA Restore stripping buffer and reprobed with other antibodies to assess signal under exactly the same loading conditions. Exposed films were scanned and the western signal for each marker (TIFF files) was quantitated using the densitometric analysis in Image J.

### Whole Mount Imaging of Fly Wings

For adult wing vein analysis, wings were dissected and kept in 100% ethanol overnight, mounted on slides in 80% glycerol in phosphate buffered saline solution, and imaged by regular light microscopy using Leica DM5500 Q microscope.

## Supplementary Figures

**Supplementary Figure 1 (for Figure 1, 2).**
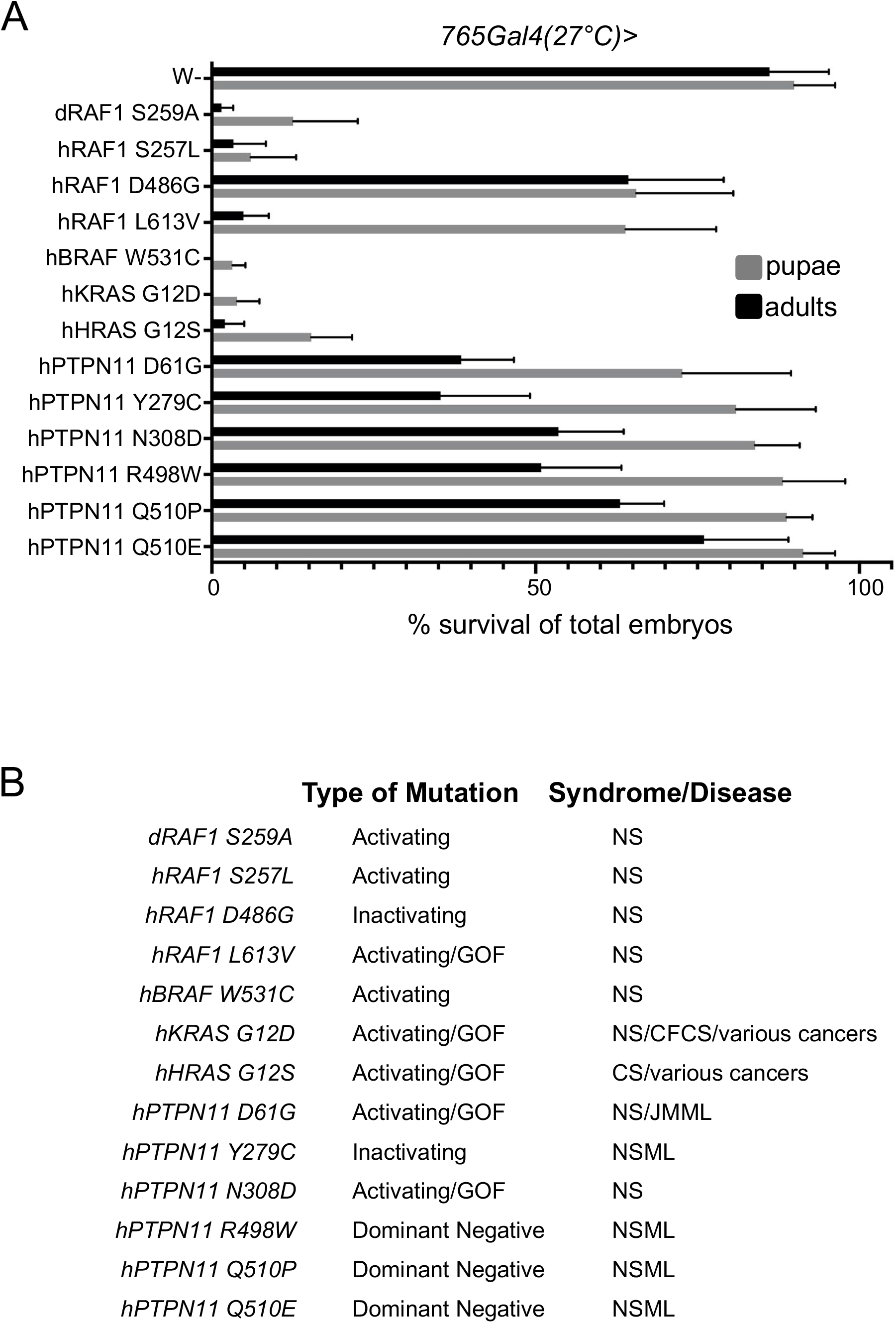
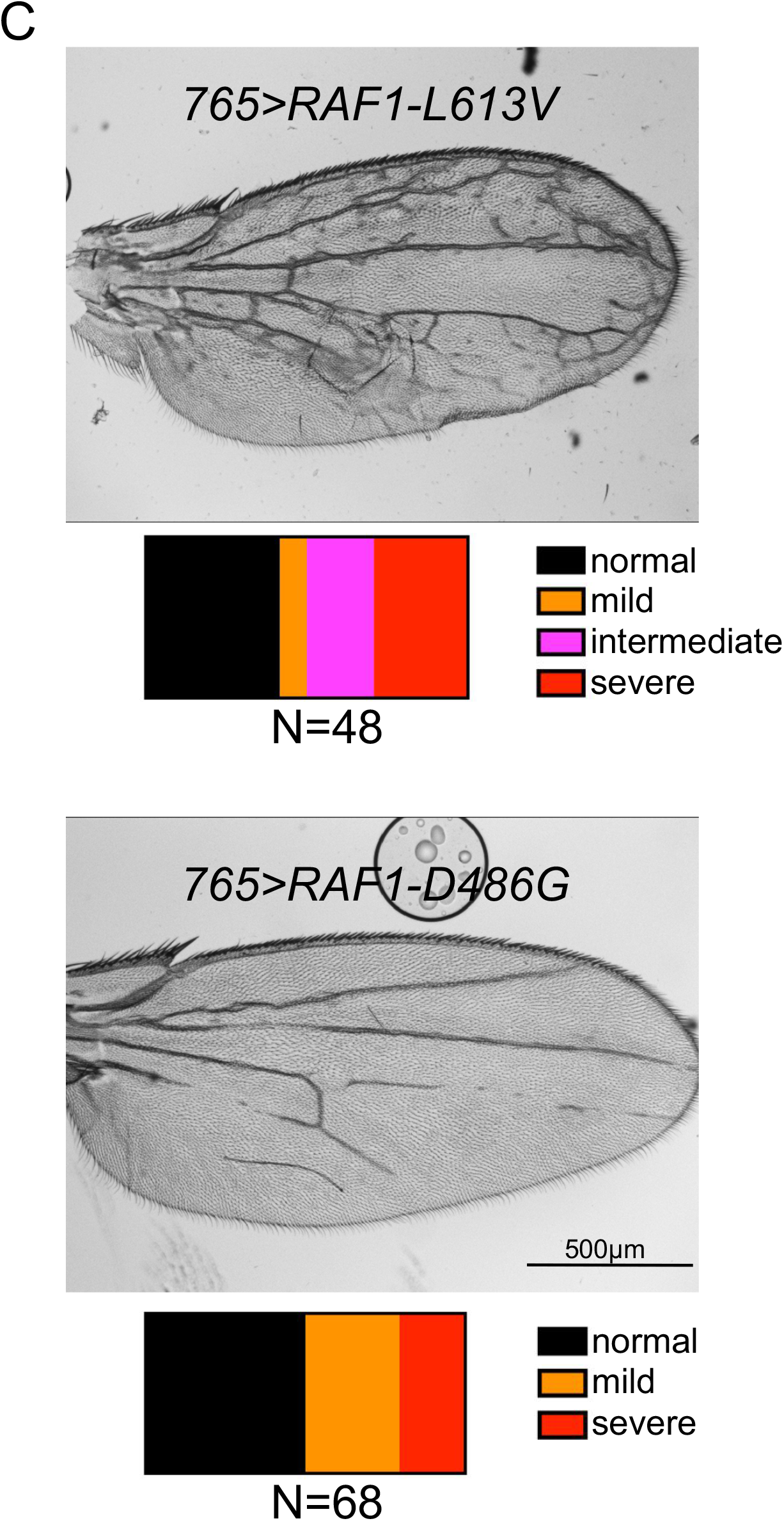
(A) Viability assay results for each RASopathy model at 27 °C. For each condition four replicates were analyzed and error bars represent standard error of the mean (SEM) here and in subsequent figures; see Methods. The percentage of surviving pupa are shown as grey bars and adults as black bars. (B) List of RASopathy variants and fly models developed in this study, the nature of the mutations, and the syndromes/diseases they are associated with in patients (Chan et al., 2008; Hijikata et al., 2017; Tartaglia and Gelb, 2010, 2005; Zheng et al., 2018). JMML-Juvenile myelomonocytic leukemia; NS – Noonan syndrome; NSML-Noonan syndrome with multiple lentigens; CFCS – cardiocutaneous syndrome; CS – Costello syndrome. (C) Bright field images of adult fly wings in which RASopathy isoforms *RAF1*^*L613V*^ and *RAF1*^*D486G*^ were expressed uniformly throughout the wing epithelia using *765-GAL4*. After adults eclosed, their wings were analyzed and the severity of wing defects assessed and binned into four categories as indicated. Finally, the proportion of each phenotype was assessed by analyzing the indicated number of wings and is represented as a color-coded bar. Similar analysis was performed with all RASopathy isoforms in Supplementary Figures 2-5. *RAF1*^*L613V*^ expression led to ectopic wing veins across the entire wing, while *RAF1*^*D486G*^ expression suppressed wing vein formation consistent with nature of the mutation inactivating the catalytic activity of RAF1.

**Supplementary Figure 2 (for Figure 2).**
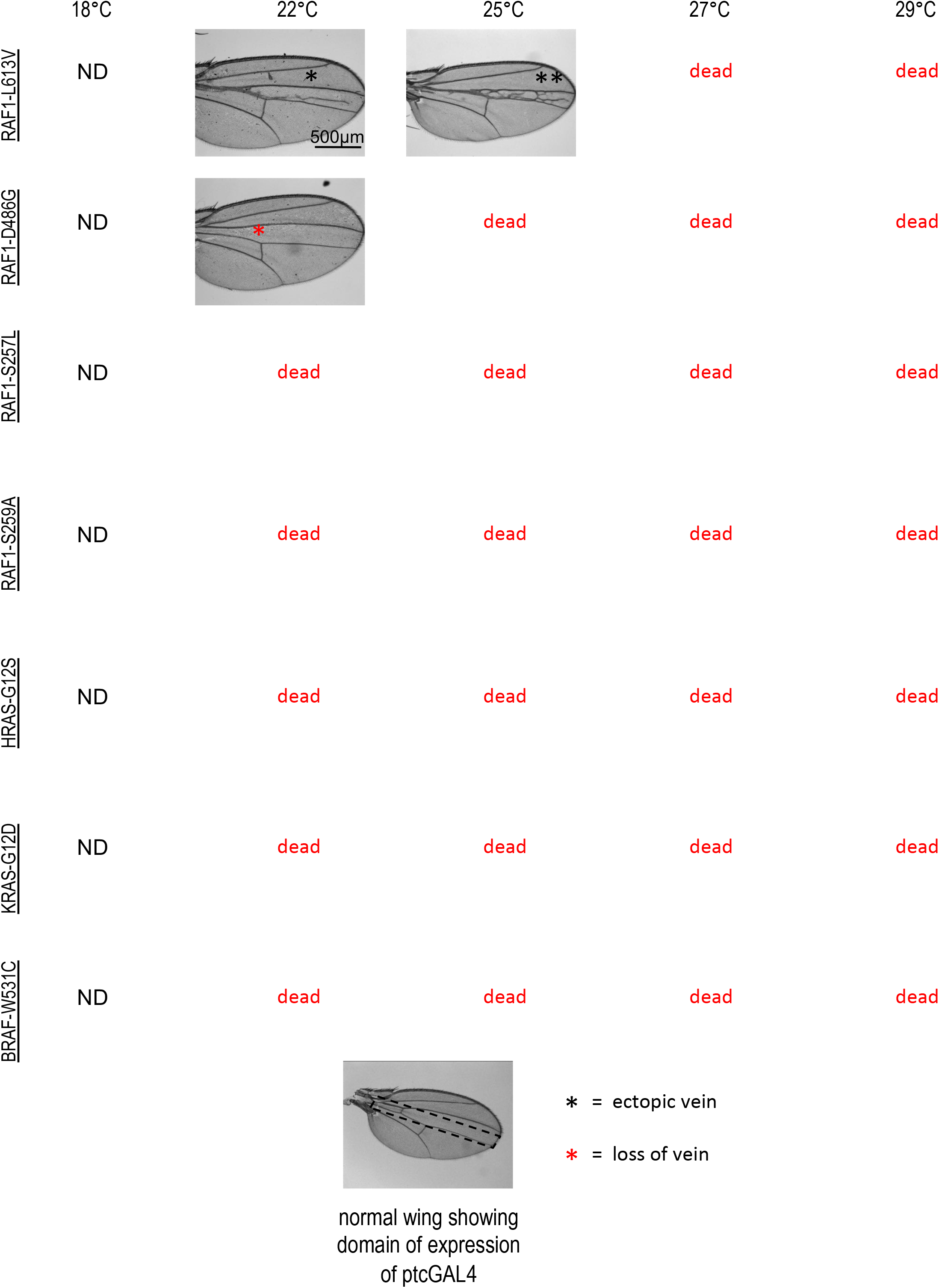
Bright field images of adult fly wings in which RAS/RAF RASopathy isoforms were overexpressed using the *ptc-GAL4* driver. The control wing at the bottom includes a dotted outline indicating the region within which *ptc-GAL4* is active. GAL4 activity progressively increases at higher temperature, leading to increased transgene expression and stronger lethality and venation phenotypes. *ptc*>*RASopathy* embryos were collected and grown at indicated temperatures; if adults eclosed, then their wings were analyzed. Black asterisk indicates ectopic veins and red asterisk indicates loss/suppression of normal veins.

**Supplementary Figure 3 (for Figure 2).**
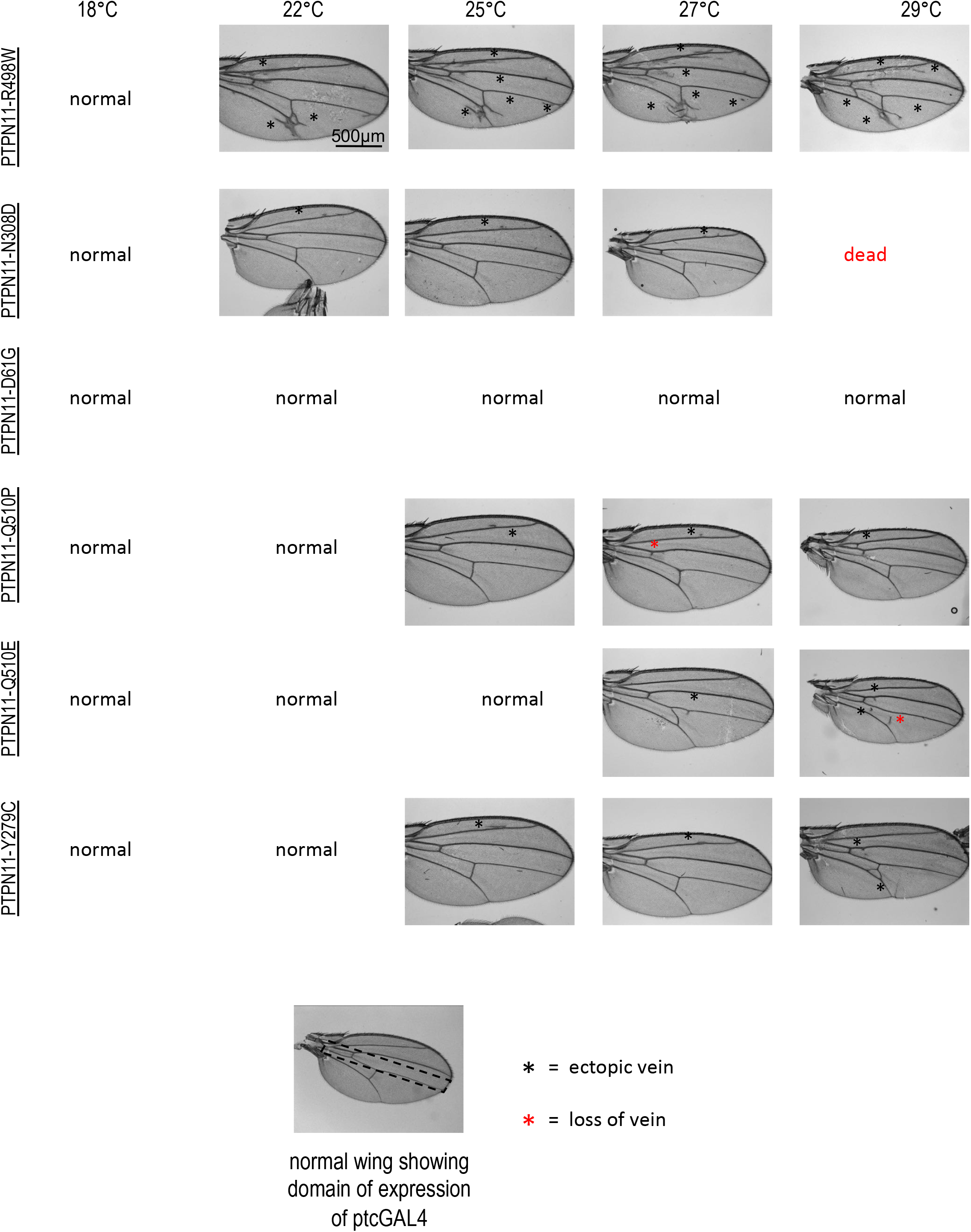
Bright field images of adult fly wings in which human *PTPN11* RASopathy isoforms were overexpressed using the *ptc-GAL4* driver. The control wing at the bottom includes a dotted outline indicating the region within which *ptc-GAL4* is active. GAL4 activity progressively increases at higher temperature, leading to increased transgene expression and stronger lethality and venation phenotypes. *ptc*>*RASopathy* embryos were collected and grown at indicated temperatures and, if adults eclosed, then their wings were analyzed. Black asterisk indicates ectopic veins and red asterisk indicates loss/suppression of normal veins.

**Supplementary Figure 4 (for Figure 2).**
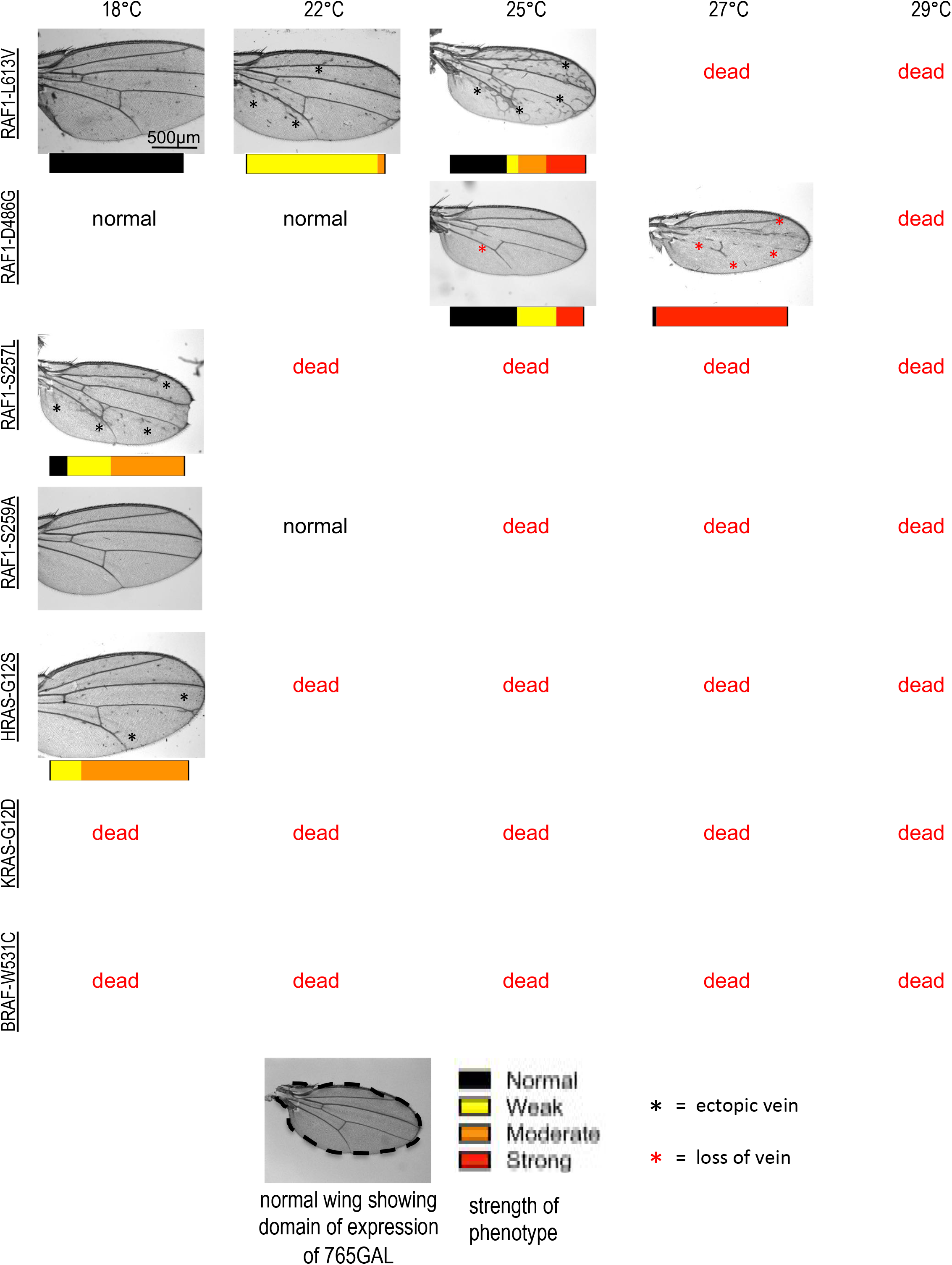
Bright field images of adult fly wings in which RAS/RAF RASopathy isoforms were overexpressed using *765-GAL4* driver. The control wing at the bottom includes a dotted outline indicating the region within which *ptc-GAL4* is active. GAL4 activity progressively increases at higher temperature, leading to increased transgene expression and stronger lethality and venation phenotypes. *765*>*RASopathy* embryos were collected and grown at indicated temperatures and, if adults eclosed, then their wings were analyzed. Penetrance and proportion of phenotypes indicated by color coded bar below some experiments. Black asterisk indicates ectopic veins and red asterisk indicates loss/suppression of normal veins.

**Supplementary Figure 5 (for Figure 2).**
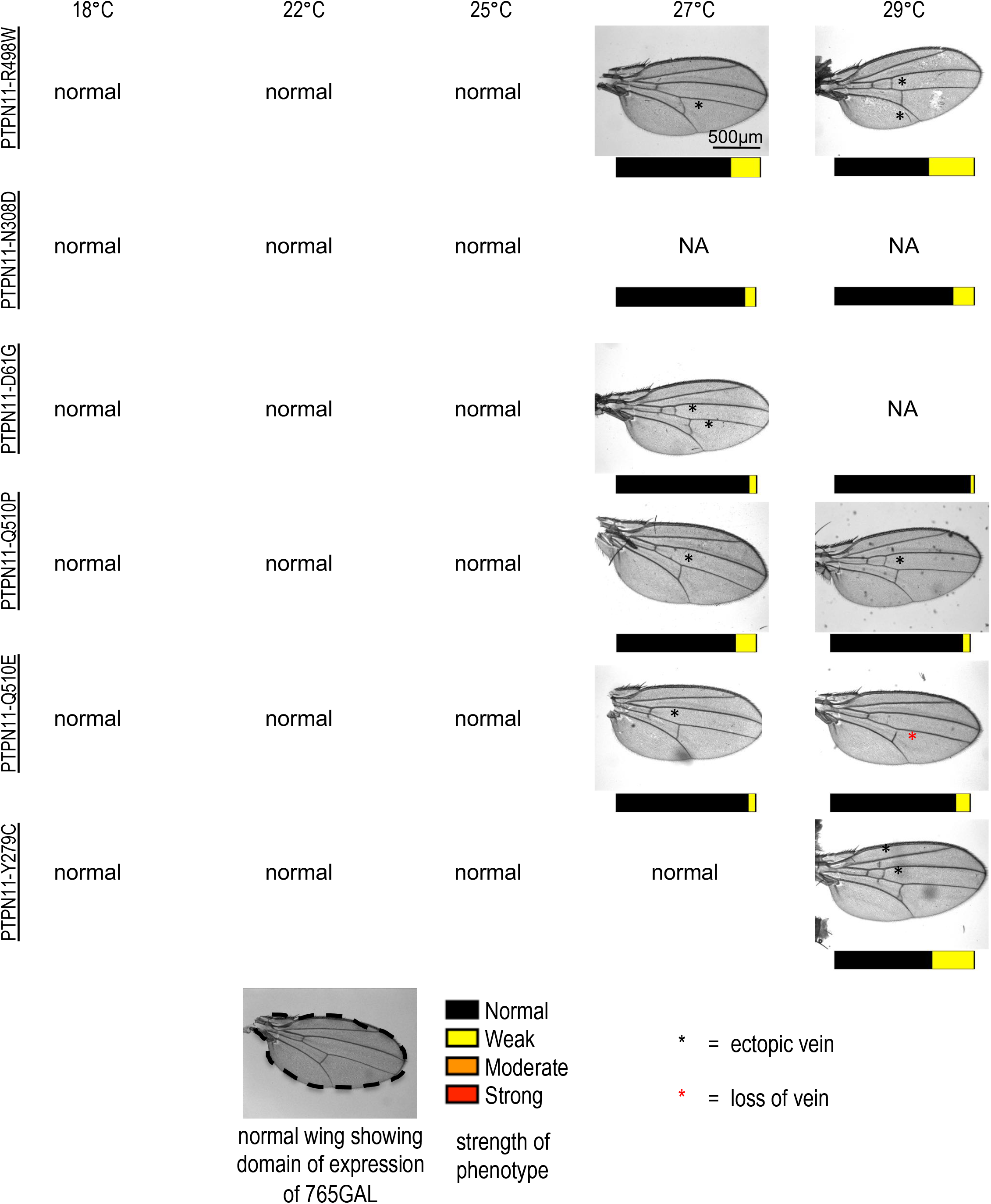
Bright field images of adult fly wings in which *PTPN11* RASopathy isoforms were overexpressed using the *765-GAL4* driver. The control wing at the bottom includes a dotted outline indicating the region within which *ptc-GAL4* is active. GAL4 activity progressively increases at higher temperature, leading to increased transgene expression and stronger lethality and venation phenotypes. *765*>*Rasopathy* embryos were collected and grown at indicated temperatures and, if adults eclosed, then their wings were analyzed. Penetrance and proportion of phenotypes indicated by color coded bar below some experiments. Black asterisk indicates ectopic veins and red asterisk indicates loss/suppression of normal veins.

**Supplementary Figure 6 (for Figures 3, 4, 5).**
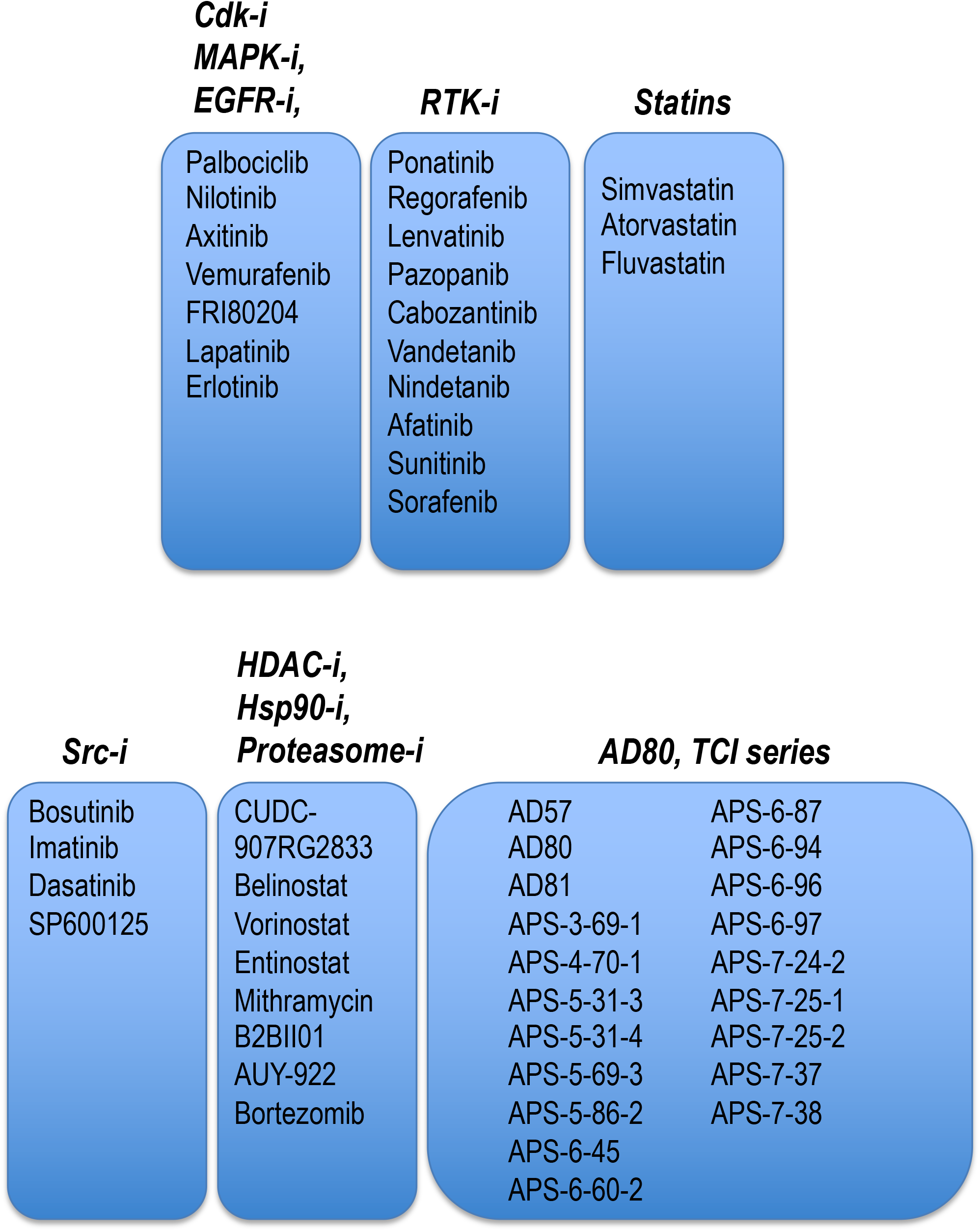
List of drugs used to screen the different RASopathy models for increased pupa and adult viability using *765-GAL4* at 27 °C as described in Figure 4. A subset of drugs that showed high efficacy in the viability screen were used for analysis of pathway activation and signaling analysis using *tubulin-GAL4*;*gal80*^*ts*^ at 27 °C as described in Figures 4, 5.

**Supplementary Figure 7 (for Figure 5).**
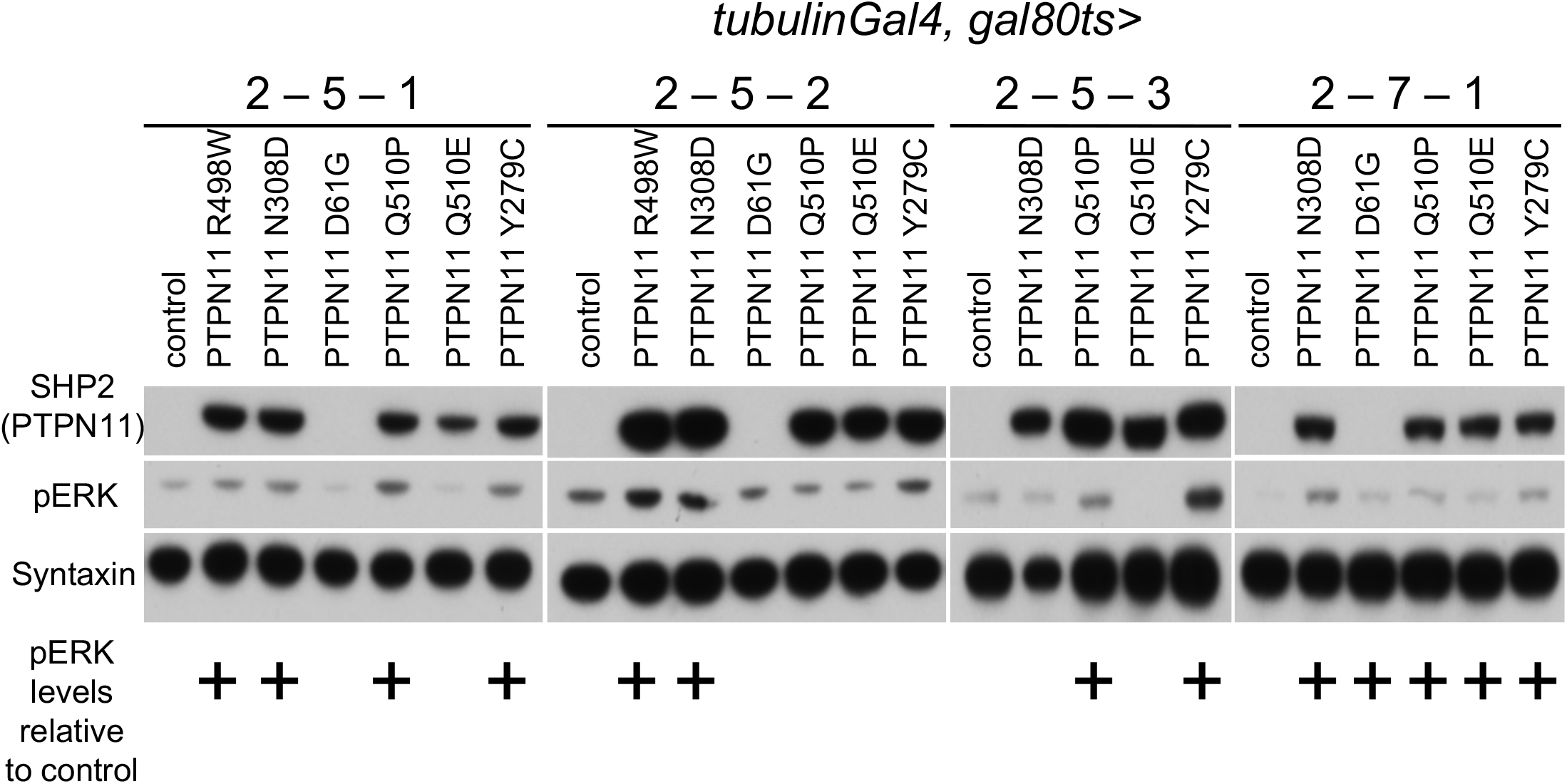
(A) Flowchart depicting timeline to induce expression of RASopathy isoforms in developing Drosophila larvae followed by western blot analysis. Embryos from flies were collected in a fixed timespan (Collection), and larva were allowed to develop at 18 °C until L3 stage (Growth). At this temperature transgene expression was not induced: basal expression of an included temperature sensitive GAL80-variant (GAL80^ts^) inhibited GAL4-dependent UAS-transgene activation. After reaching L3 stage the larvae were shifted to 27 °C, which led to destabilization of GAL80^ts^ protein and induced expression of the RASopathy-encoding transgenes (Induction). After a fixed time of induction, larvae were collected and whole-body lysates extracted for western blot analysis. (B) Western blot analysis of different growth, collection, and induction conditions (C-G-I) for analysis of pathway activation by *PTPN11* isoforms. Whole larval lysates were collected from different *PTPN11* lines under the indicated conditions. Western blot analysis performed to detect MAPK pathway activation. Some isoforms upregulated pERK levels in the 2-5-1 condition but downregulated it in the 2-5-2 condition (−Q510P and −Y279C). This suggests feedback-induced downregulation of upstream MAPK components such as phosphorylated ERK (pERK), a common feature of the MAPK pathway. Other isoforms also show complex regulation of pERK but all isoforms increase pERK compared to control in the 2-7-1 condition. The SHP2 (*PTPN11*) antibody detected mostly equivalent levels of expression across *PTPN11* lines. −D61G was not detected using the SHP2 antibody, perhaps due to mutation-induced loss of the antibody epitope; its presence was confirmed by genomic PCR of fly models.

**Supplementary Figure 8 (for Figure 6).**
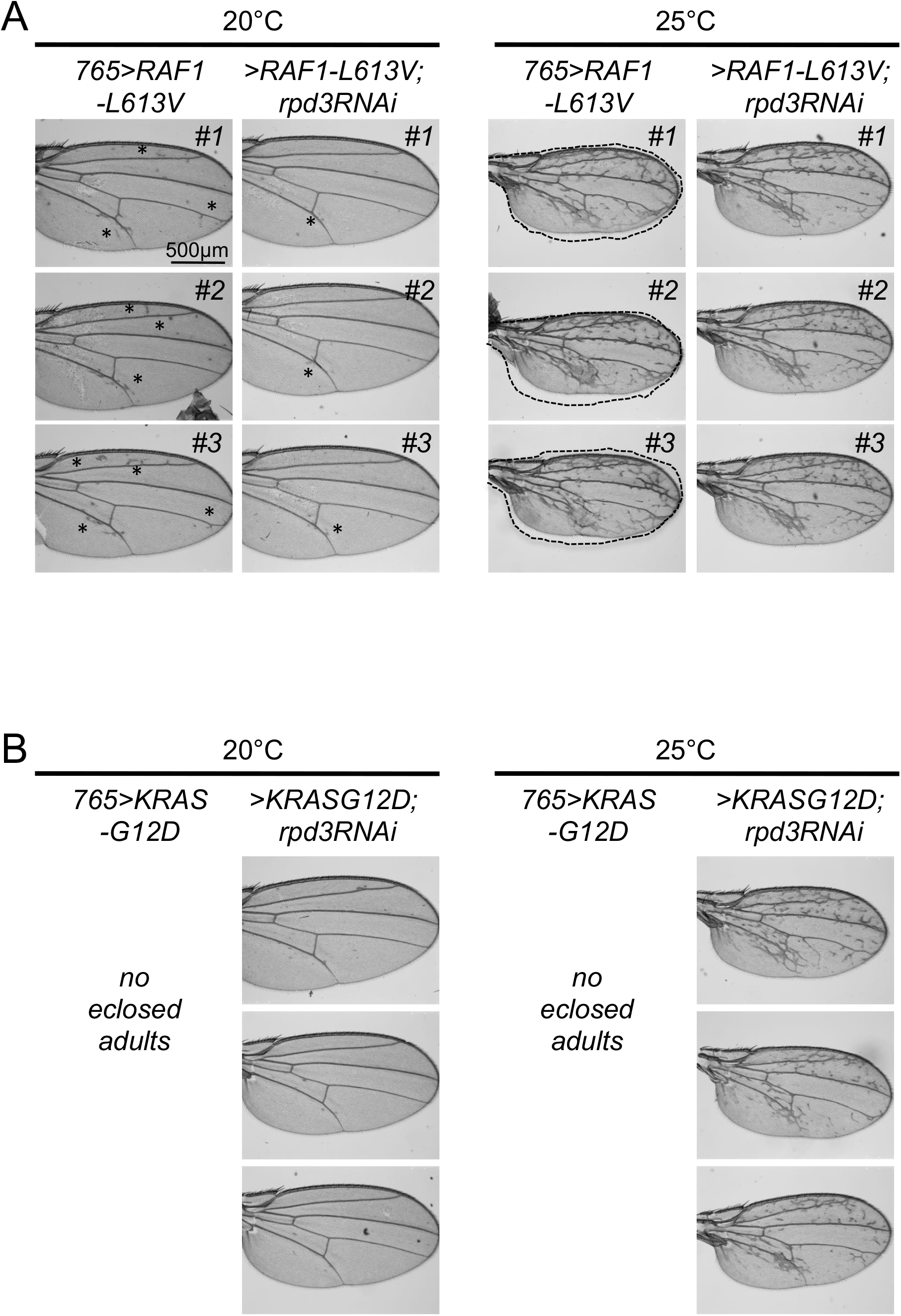
(A) Genetic modifier experiments with *RAF1*^*L613V*^ isoforms show dependency on HDAC1. Each isoform was expressed throughout the developing larval wing disc using the *765-GAL4* driver under different temperature conditions. *765>RAF1*^*L613V*^ flies exhibited ectopic wing venation phenotypes 20 °C, which was suppressed by RNAi-mediated knockdown of the fly HDAC1 ortholog Rpd3 (*765*>*RAF1*^*L613V*^,*rpd3-RNAi*). The suppression of ectopic wing venation did not occur at 25 °C with stronger induction of the isoform; interestingly at 25 °C, knockdown of Rpd3 consistently altered wing size as shown by dotted outline of the *765*>*RAF1*^*L613V*^,*rpd3-RNAi* wing superimposed on *765>RAF1*^*L613V*^ wing images. Black asterisk indicates ectopic veins. Three wings of each experiment are shown for comparison. (B) Genetic modifier experiments with the *KRAS*^*G12D*^ isoform demonstrated functional dependency on HDAC1 for ectopic wing venation. When *765>KRAS*^*G12D*^ flies were raised at 20 °C and 25 °C no adults eclose; this developmental lethality was suppressed by knockdown of *Rpd3*, resulting in adult eclosure. At 20 °C, *765*>*KRAS^G12D^,rpd3-RNAi* flies exhibited near-normal wing vein patterning. At 25 °C the ectopic wing venation pattern was not suppressed, again presumably due to stronger induction of the *KRAS*^*G12D*^ transgene. Three wings for each experiment are shown for comparison.

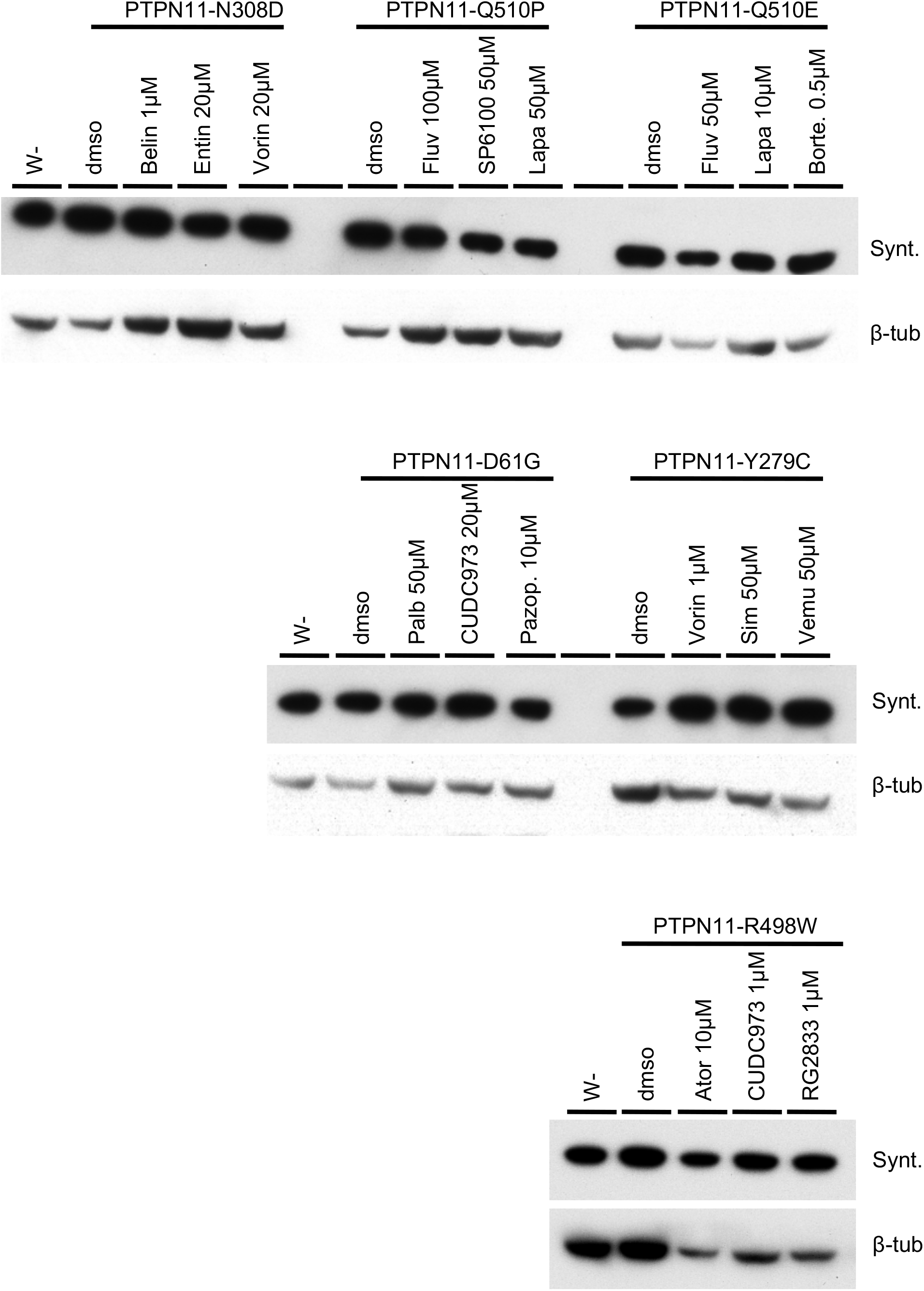

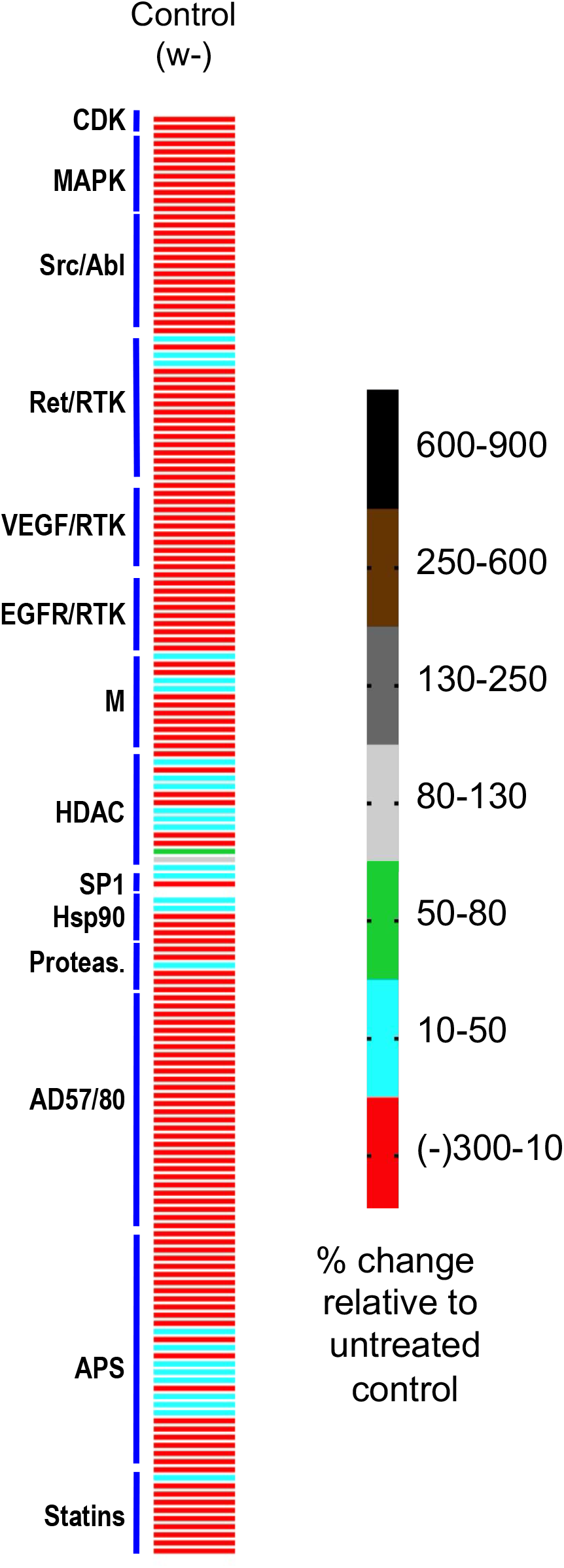

